# Actinobacteriophage Inteins: Host Diversity, Local Dissemination, and Non-Canonical Architecture

**DOI:** 10.1101/2025.01.02.630785

**Authors:** Sophia P. Gosselin, Danielle Arsenault, Johann Peter Gogarten

## Abstract

Intein presence within Actinobacteriophages (within PhagesDB) was last surveyed in 2016, and despite a 5-fold increase in the size of the database, has not been updated since. To address this, we present a modern survey of the current iteration of the PhagesDB database. We developed a new algorithm — Iterative Cluster Expansion BLAST (ICE-BLAST) — to expand our search to more divergent sequences. Nearly 800 inteins were retrieved through this process; the majority of which were previously unreported. We describe the nature of these inteins, their classes, integration target sites, distribution within phage clusters, and explore the geographical location of nearly identical intein sequences found in divergent exteins. Our findings suggest that these inteins recently invaded local phage populations. We also find two instances of a Cas4 exonuclease intein evolving from a terminase large subunit intein, and propose a model by which one of these inteins was able to utilize sequence similarity conferred by a shared nucleotide binding site to jump between genes. Additionally, we find inteins with never-before-reported homing endonucleases, and inteins with homing endonucleases encoded in a reading frame separate from that which encodes the extein and the intein’s self-splicing domain. We provide predicted structures for these elements and hypothesize on their evolution and relation to free-standing homing endonucleases within phage genomes. Finally, we provide evidence that these “non-canonical” inteins are still transferring between host genomes, in a fashion similar to other inteins with canonical homing endonucleases within the dataset.

## Introduction

Inteins are self-splicing, site-specific, mobile genetic elements that invade protein-encoding genes across all domains of life and viruses, including bacterial viruses (phages) (Gogarten et al. 2002; Lennon and Belfort 2017; Nanda et al. 2020). An intein begins its journey as an intervening DNA sequence at a specific insertion site within the coding sequence of a host gene. Unlike the intervening DNA sequences known as introns which are post-transcriptionally removed at the RNA level, an intein is transcribed and translated together with its host gene. The intein then seamlessly excises itself post-translationally from the host protein, referred to as the extein, through the unique process of protein splicing (Shih et al. 1988; Hirata et al. 1990; Kane et al. 1990). Protein splicing, performed by the self-splicing domain of the intein, yields a functional extein and the free intein. Inteins are divided into three classes depending on their initial steps of splicing, the residues involved, and differences in reaction chemistry (Tori et al. 2010; Nanda et al. 2020).

Once excised from the extein, the intein uses its homing endonuclease (HEN) domain to engage in the self-propagation method known as homing (Goddard and Burt 1999). When in the presence of a copy of its host gene lacking the intein’s DNA sequence at its particular insertion site, the HEN makes a double-strand break at the intein insertion site. Through homologous recombination-based repair using the intein-containing copy of the gene as the template, the intein sequence is copied into the previously vacant insertion site (Belfort and Roberts 1997). Homing allows inteins to rapidly rise to saturation in populations, largely through horizontal transfer-based homing, at which point there is no longer selective pressure to maintain the HEN (Goddard and Burt 1999). The HEN degrades, turning the full intein into a mini intein which contains only the self-splicing domain (Iwaï et al. 2017). Sometimes, an intein may acquire an internal stop codon in the HEN as it degrades, generating a split intein (Tavassoli and Benkovic 2007; Dassa et al. 2009; Hoffmann et al. 2021). Eventually, the mini and split inteins are lost and the intein-free allele once again dominates. Newer models propose the co-existence of full, mini, split, and empty alleles as opposed to synchronized progression through the phases (Gogarten and Hilario 2006; Yahara et al. 2009; Barzel et al. 2011). As an alternate evolutionary endpoint to degradation and loss, some inteins have also been inferred to have acquired novel functions, acting as regulatory elements under stressful conditions (Novikova et al. 2016; Lennon et al. 2018) or evolving into mating-type switching endonucleases such as HO of *Saccharomyces cerevisiae* (Coughlan et al. 2020).

Several different HEN domain types have been reported in inteins. Most inteins possess canonical LAGLIDADG homing endonucleases, which are in-frame with the rest of the intein (Gorbalenya 1998; Gimble 2000). Others include: inteins with an in-frame HNH endonuclease (Dalgaard, Klar, et al. 1997), split-inteins possessing VSR-like endonucleases (Dassa et al. 2009), and LAGLIDADG HENs encoded on the anti-sense strand (Gorbalenya 1998; Gogarten et al. 2002). Interestingly, most of these non-canonical inteins are found within phage genomes, with actinobacteriophages serving as a particular hot-spot in research.

Actinobacteriophages are promising therapeutic alternatives to antibiotics in the treatment of multi-drug resistant actinobacterial infections (Allué-Guardia et al. 2021; Zeynali kelishomi et al. 2022; Dedrick et al. 2023; Yang et al. 2024). However, much is still unknown about actinobacteriophage structure, function, and genomic evolution, necessitating increased studies in these areas. In recent years, teaching programs (SEA-PHAGES and PHIRE) employing citizen-science based approaches to isolate and sequence actinobacteriophages have added thousands of genomes to databases such as PhagesDB (Russell and Hatfull 2017). PhagesDB thus serves as an ideal candidate for large-scale computational analyses aiming to dissect phage genomic evolutionary dynamics, with inteins serving as particularly powerful tools for tracing genetic exchange between phage populations (Soucy et al. 2014).

Inteins are present within actinobacteriophages (Tori et al. 2010; Tori and Perler 2011; Kelley et al. 2016). However, since the last survey of these inteins (Kelley et al. 2016) the PhagesDB database has expanded from 841 to 5075 genomes; providing a unique resource to study the distribution and evolution of phage inteins. Hence, we conducted a new survey of PhagesDB, accessed in September of 2023. To capture more divergent inteins within the database, we developed an algorithm, ICE-BLAST, which uses a combination of PSI-BLAST and USEARCH clustering to iteratively search an amino acid database using the representative sequences of clusters (centroids) of matches from previous searches. Herein we discuss the development and testing of this algorithm, and the inteins discovered through its application to PhagesDB. We investigate intein distribution within phage clusters, variation in host genes, and geographic dissemination. Through the latter, we found evidence supporting events of rapid geographically localized propagation of intein alleles into new populations. Our phylogenetic analyses also uncovered two occurrences of class 3 inteins of terminase large subunits sharing a common ancestor with Cas 4 exonuclease inteins. Additionally, we report several cases of inteins with HEN domains and architectures that have rarely been reported on, or were previously unseen. These include HENs encoded within a separate ORF on the same strand as the intein, and HENs in a separate ORF on the anti-sense strand.

## Results

### Intein Distribution across Actinobacteriophage Clusters

Using our program ICE-BLAST, tested using simulated data from our program Invader-Sim, we identified 784 putative intein sequences in PhagesDB (details on the intein retrieval and validation process are provided under Materials and Methods). A table containing information on all 784 inteins including their host phage, host protein (extein), splicing class, and HEN type is available (Supplemental Table 1). These 784 inteins span a wide variety of PhagesDB phage clusters (Figure 1), with the majority being in clusters C1 (415), A1 (92), and E (68), all of which are listed as mycobacteriophage clusters. The other 209 inteins were distributed across 31 phage clusters and several singleton phages (phages not assigned to a cluster). Of the 31 phage clusters, the maximum number of inteins observed in one cluster was 35, and the majority of clusters contained less than 10 inteins. The most intein-rich genomes were all from cluster C1 phages, with phage Cane17 having seven inteins, followed by phages Blubba and Colt with six inteins each.

**Figure 1.**
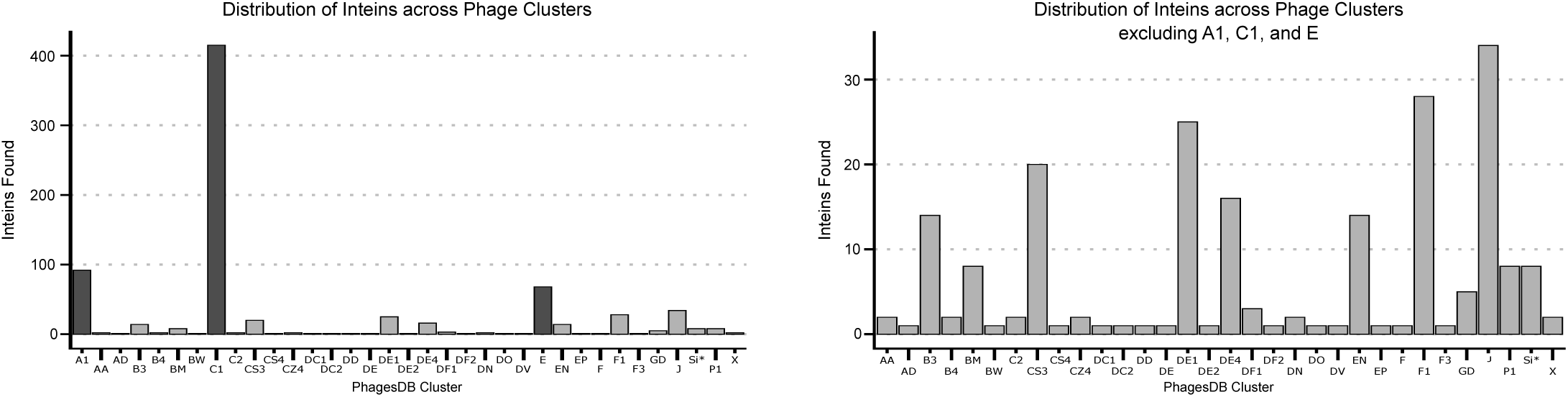
Total number of inteins found in each PhagesDB phage cluster, depicted with (left) and without (right) intein-rich clusters A1, C1, and E. A1, C1, and E are depicted with dark gray bars and contained 92, 415, and 68 inteins respectively. Singletons are denoted as Si*.

### Wide Range of Actinobacteriophage Proteins Invaded by Inteins

By re-annotating the host protein sequences of the 784 putative inteins, we found that they invade 19 distinct proteins (Figure 2). Unsurprisingly, the majority were found in proteins previously reported as containing inteins. These include: Cas4 exonuclease (InterPro IPR013343), DNA helicase (InterPro IPR006500), DNA methylase N-6/N-4 (Pfam PF01555) and C-5 (InterPro IPR001525), DNA primase (InterPro IPR006171), nucleotidyltransferase (InterPro IPR018775), phosphoesterase (InterPro IPR029052), portal protein (InterPro IPR009279), terminase large subunit (Pfam PF04466), and thymidylate synthase (InterPro IPR003669). The remaining host proteins are new to this study within actinobacteriophages, and include: DNA polymerase III alpha subunit (InterPro IPR011708), minor capsid protein (InterPro IPR006528), ribonucleotide reductase subunits alpha (InterPro IPR026459) and beta (InterPro IPR026494), RtcB-like RNA ligase (InterPro IPR021122), tape measure protein (InterPro IPR013491), and tRNA splicing RNA ligase (InterPro IPR036025). Two types of intein-containing exteins identified could not be definitively annotated, but have best matches to baseplate wedge domain containing proteins (UniProt M9MUF4) and integrin-like proteins respectively. Based on the annotations of other closely related intein-containing exteins, the former could be a portal protein of some sort, but does not share considerable similarity to known portal proteins in any NCBI database. The inteins present within the putative integrin-like exteins were previously reported on in Kelley et al. 2016, but with fewer intein-containing exteins (Kelley et al. 2016).

**Figure 2.**
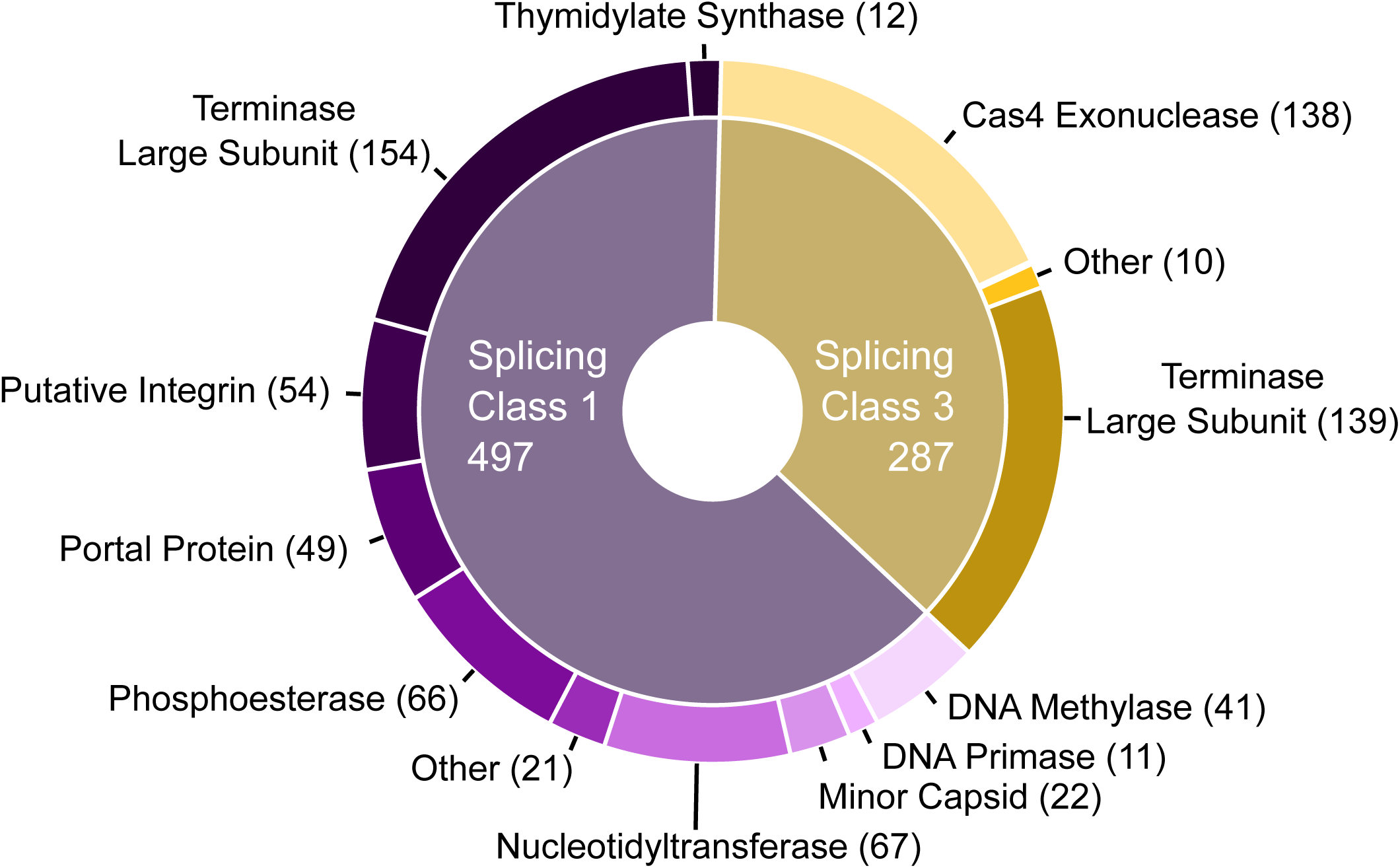
Identified inteins, which fall into class 1 and class 3 based on their splicing domains, are distributed across 19 different host genes: baseplate wedge domain protein (InterPro M9MUF4), Cas4 exonuclease (InterPro IPR013343), DNA helicase (InterPro IPR006500), DNA methylase N-6/N-4 (Pfam PF01555) and C-5 (InterPro IPR001525), DNA polymerase III alpha subunit (InterPro IPR011708), DNA primase DnaG-like (InterPro IPR006171), putative integrin-like protein, minor capsid protein (InterPro IPR006528), nucleotidyltransferase DNA polymerase beta superfamily (InterPro IPR018775), phosphoesterase (InterPro IPR029052), portal protein (InterPro IPR009279) ribonucleotide reductase alpha subunit (InterPro PR026459) and beta subunit (InterPro IPR026494), RtcB-like RNA ligase (InterPro IPR021122), tape measure protein (InterPro IPR013491), terminase large subunit (Pfam PF04466), thymidylate synthase (InterPro IPR003669), and tRNA-splicing ligase (InterPro IPR036025).

### Variation in Intein Splicing Domains

The putative inteins were clustered at 70% sequence identity. For each intein cluster, we determined the intein splicing class (1, 2, or 3). Most of the inteins were class 1 (497) while the remainder (287) were class 3. The class 1 inteins displayed a much wider range of host proteins (16 different proteins) than the class 3 inteins which were only found within 4 different extein hosts. More than half of the inteins within our dataset (431) invaded just two proteins, with 293 inteins within terminase large subunit sequences, and 138 within Cas4 exonucleases (Figure 2). The size of class 1 inteins ranged from 385 amino acids (aa) (terminase large subunit of cluster DC2 phage Clown) to 134 aa (terminase large subunit of cluster EN phage C3PO). The size of class 3 inteins ranged from 359 aa (DNA helicase of cluster C1 phage Alice) to 225 aa (minor capsid of singleton phage REQ3). In addition to classifying the splicing domain of each intein, we classified their HEN domains, which revealed further diversity and previously undocumented intein architectures.

### Variation in Intein Homing Endonuclease (HEN) Domains

Most of the inteins in our database (712) possess classic intein architecture: a self-splicing domain and a canonical LAGLIDADG (InterPro IPR004860) HEN domain. Of the remaining 72 inteins, nine have decaying or completely degraded HENs (mini inteins), and 63 possess unique non-canonical HENs. These non-canonical HENs include in-frame HNH (InterPro IPR002711) HENs (Figure 3A), LAGLIDADG or VSR-like (InterPro IPR004603) HENs encoded on the anti-sense strand (Figure 3B-C), and out-of-frame endonuclease VII-like (InterPro IPR004211) HENs (Figure 3D). The inteins possessing these non-canonical HENs are distributed across four host protein types: DNA polymerase, minor capsid, terminase large subunit, and thymidylate synthase (Figure 3E). Based on current knowledge, these four non-canonical HEN architectures are rare, with some being reported on for the first time with this work.

**Figure 3.**
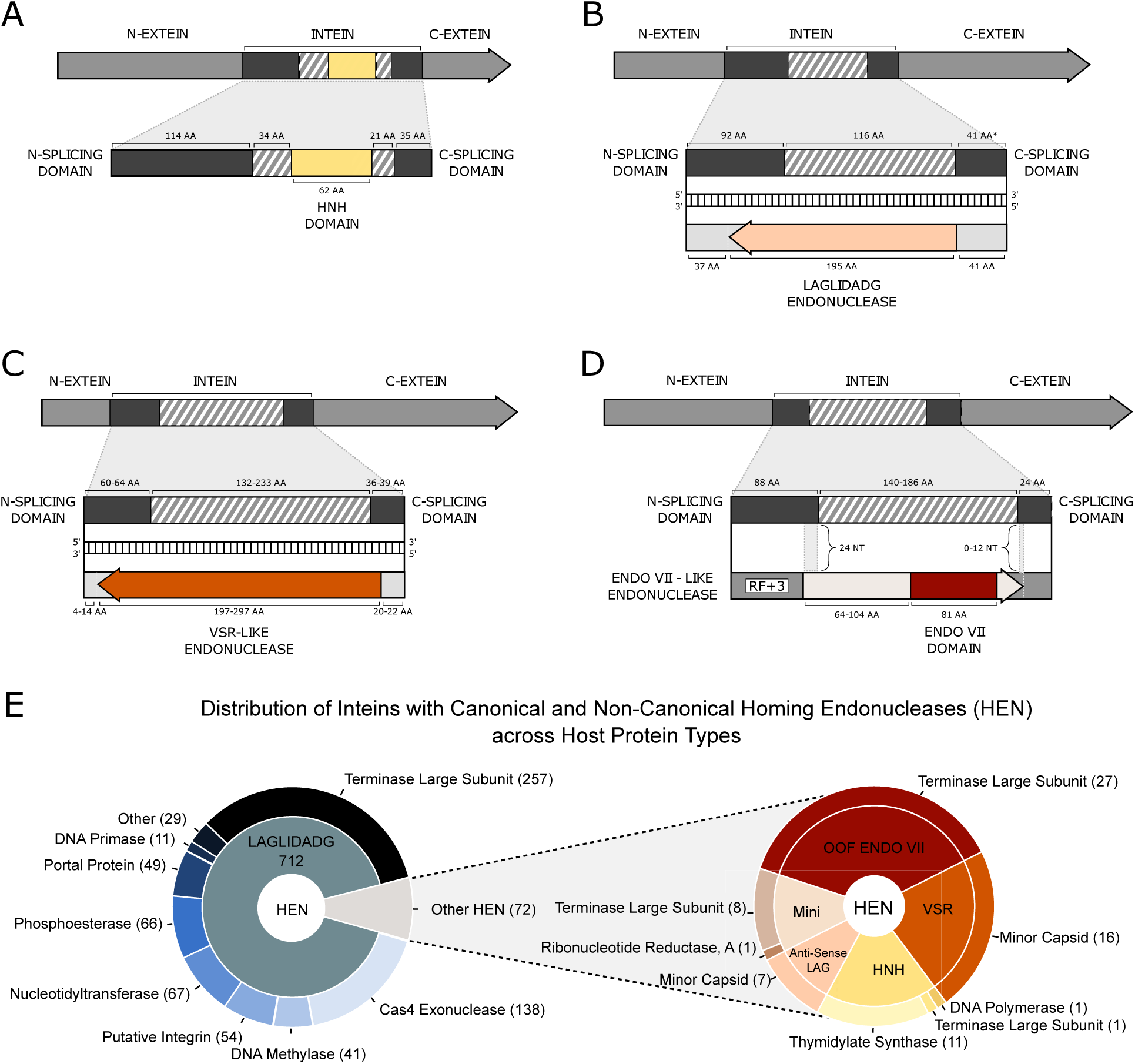
A-D. Gene-map depictions of the four observed non-canonical intein HEN organizations. Regions shaded with diagonal lines indicate HEN-encoding region. **A.** in-frame HNH HEN. **B.** anti-sense LAGLIDADG HEN. **C.** anti-sense VSR-like HEN. **D.** endonuclease VII-like HEN in reading frame 3 relative to intein. **E.** Distribution of inteins with canonical and non-canonical HEN architectures across host protein types.

The non-canonical HENs which have already been reported on include the in-frame HNH HEN which is observed in phages (Dalgaard, Klar, et al. 1997; Dalgaard, Moser, et al. 1997), the anti-sense LAGLIDADG HENs (Gorbalenya 1998; Gogarten et al. 2002), and the VSR-like HEN which was reported in the context of split inteins with no other known intein-associated reports (Dassa et al. 2009). The fourth type of non-canonical HEN we found, an endonuclease VII-like HEN encoded within the HEN region but in frame 3 relative to the rest of the intein, has not yet been reported to our knowledge.

To further assess the identities of the out-of-frame and anti-sense non-canonical HENs, AlphaFold-predicted structures of the inteins (Supplemental Figure 1) and the HENs in their respective frames (Supplemental Figure 2) were compared to solved structures. Based on the predicted structures for the endonuclease VII-like and anti-sense LAGLIDADG HENs, coupled with the fact that these HENs can be dimeric, the terminal domains of these structures may be linker domains or secondary DNA binding domains (Raaijmakers et al. 1999; McMurrough et al. 2018). The VSR-like HEN appears to contain a long stretch of linked turns and alpha helices not present in normal VSR endonuclease structures (Hennecke et al. 1991). The predicted local distance difference test (pLDDT) — a per-residue measure of local prediction confidence (Jumper et al. 2021) — for this stretch is relatively high, especially within helices, but the region lacked meaningful matches against PDB structures. However, an HHPred search found a significant match (e-value of 0.000046) against protein structure 3R3P from Bacillus phage 0305phi8-36, which indicates the VSR-like HEN may be an EDxHD HEN (Taylor et al. 2011).

### Actinobacteriophage Intein Phylogenetics

Having characterized the structural architectures of each intein, we sought to investigate how these inteins have evolved with respect to their host proteins and host phages. Performing such evolutionary characterizations of inteins can often unveil pathways for genetic exchange within populations that may have gone unnoticed when focusing on genetic material with primarily orthodox vertical inheritance (Soucy et al. 2014; Turgeman-Grott et al. 2023). Thus, we constructed phylogenies from the DNA sequences of all inteins in our database. The extracted subsets of intein sequences used to generate these phylogenies were separated by HEN type, with the largest set (canonical HENs) split into two sets based on splicing class (class 1 or class 3). By analyzing the grouping patterns of inteins within the phylogenies compared to the types of host proteins they invade, potential instances of intein host protein switching (e.g. a terminase large subunit-targeting intein evolving into a Cas4 exonuclease-targeting intein) can be identified.

While some inferences could be drawn from the phylogenies of the inteins with non-canonical HEN domains, current sampling is ultimately too sparse to make any definite claims from the intein phylogenies alone (Supplemental Figures 3-6). In contrast, the inteins with canonical HENs produced more informative phylogenies.

The class 1 inteins with a canonical LAGLIDADG HEN produced a phylogeny which provided evidence for several independent events of intein integration site diversification events (Figure 4). For example, the terminase large subunit inteins form a polyphyletic clan with at least four separate groups. Each group is separated, divided from the other terminase large subunit groups by multiple highly supported splits along the inner nodes. Similarly, portal protein inteins and DNA methylase inteins form polyphyletic clans with five and three separate groups respectively.

**Figure 4.**
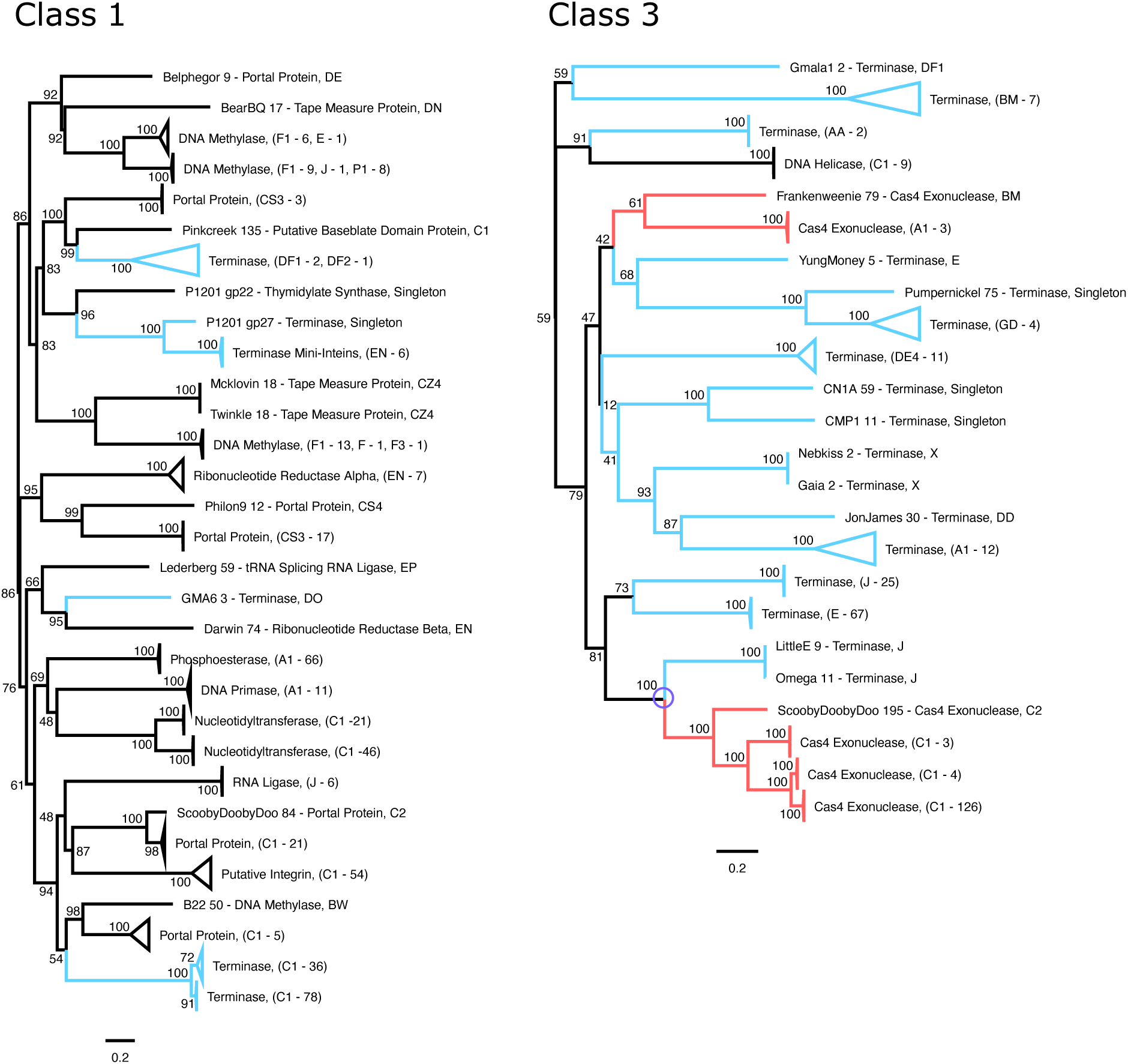
The DNA sequences of the inteins with canonical LAGLIDADG HEN domains were split into two subsets by splicing class (class 1 and class 3), with each subset then used for maximum likelihood phylogenetic reconstruction. Substitution models used were TIM2+F+R3 (class 1) and TPM2+F+R4 (class 3). Ultrafast bootstrap support is listed on internal nodes. Scale bar measures substitutions per site. Clans of inteins within the same extein are collapsed; clade labels indicate the number of inteins in the collapsed group, and which phage clusters they are found in. Branches leading to terminase large subunit inteins are colored in blue, and branches leading to Cas4 exonuclease inteins are colored in red. On the class 3 phylogeny, the potential extein switching event between terminase large subunit and Cas4 exonuclease inteins (involving phages LittleE and ScoobyDoobyDoo) further investigated is indicated with a purple circle.

The class 3 inteins with canonical LAGLIDADG HENs are almost entirely found within Cas4 exonuclease and terminase large subunit exteins (Figure 4). Within this phylogeny are two instances of Cas4 associated inteins diverging from terminase large subunit inteins. One of these groupings had particularly high bootstrap support, and was thus further investigated as a potential case of extein switching (an intein evolving to target an insertion site inside of a new type of host protein).

### Extein Switching

The class 3 intein phylogeny (Figure 4) depicts two instances of a clade of Cas 4 exonuclease inteins grouping inside of a clade of terminase large subunit inteins. This grouping suggests that an intein which now targets an insertion site in Cas4 exonucleases evolved from an intein which targets an insertion site in terminase large subunits. We will refer to this type of evolutionary event as extein switching. To assess whether this extein switching topology was a phylogenetic artifact, an approximately unbiased (AU) tree topology test (Shimodaira 2002) was used to gauge the strain induced when one attempts to force the terminase large subunit and Cas4 exonuclease inteins to group by extein type, as opposed to the observed topology. The same model determined by IQ-TREE’s ModelFinder for the original class 3 intein phylogeny (WAG+F+R4) was used for the AU test. The test produced little support for the grouping of these inteins by extein type, (bootstrap proportions of 0.998 and 0.002 for tree constrained by extein switching event and the tree constrained by extein type respectively (Supplemental Table 2)). After investigating both putative extein switching events, the event involving phages LittleE and ScoobyDoobyDoo provided a more complete picture of how the extein switching may have occurred. The terminase large subunit inteins of the other extein switching event (involving phage Frankenweenie (Figure 4)) all exhibited degradation in their HEN domains, making comparison of predicted intein structures for this group less informative. Thus, the event involving phages LittleE and ScoobyDoobyDoo will be the focus of the presented investigation that follows.

### Extein Switching from Terminase Large Subunit to Cas4 Exonuclease

Comparing the +/-30 nucleotides at the intein insertion site across the LittleE terminase large subunit and ScoobyDoobyDoo (SDD) Cas4 exonuclease reveals regions of striking similarity (Supplemental Figure 7A). Both insertion sites reside in stretches of DNA sequence which encode important nucleotide binding sites for their respective proteins (Supplemental Figure 7B). Comparison to the solved structure of a phage terminase large subunit bound to ATP (Zhao et al. 2013) shows the intein insertion site encodes a region on an alpha-helix within the Rossmann fold constituting the nucleotide binding site within the ATPase domain of the terminase large subunit. The SDD Cas4 exonuclease intein insertion site encodes the motif EFKT, which is the RecB-like motif-III in the Cas4 exonuclease, and is important for nucleotide binding (Shiimori et al. 2018). In addition to assessing insertion site sequence similarity, we compared the predicted protein structures for the inteins. Alignment of the predicted structures generated for the terminase large subunit intein of LittleE, the Cas4 exonuclease intein of SDD, and the terminase large subunit intein of cluster DE4 phage APunk (representing a clade of terminase large subunit inteins which do not group with LittleE’s terminase large subunit intein) reveals them all to be near identical, barring APunk (Figure 5). These results align with the expected results based on the phylogeny. The APunk intein does not align as strongly to the LittleE and SDD inteins as they do to each other, specifically exhibiting a slight structural difference in its HEN (Figure 5). This difference in HEN structure between the APunk and LittleE terminase large subunit inteins in particular reflects how they target two separate regions of the terminase large subunit gene (both intein insertion site regions encode alpha-helices in the ATPase domain, but two distinctly different helices). In contrast, the high similarity between the HEN domains of the SDD and LittleE predicted intein structures further supports the notion that these inteins can recognize and cut an extremely similar intein insertion site. To explain the lack of intein in SDD’s terminase large subunit, we later propose a model in which the SDD terminase large subunit was not a viable host for the LittleE terminase large subunit intein, and the SDD Cas4 exonuclease provided a viable alternate insertion site.

**Figure 5.**
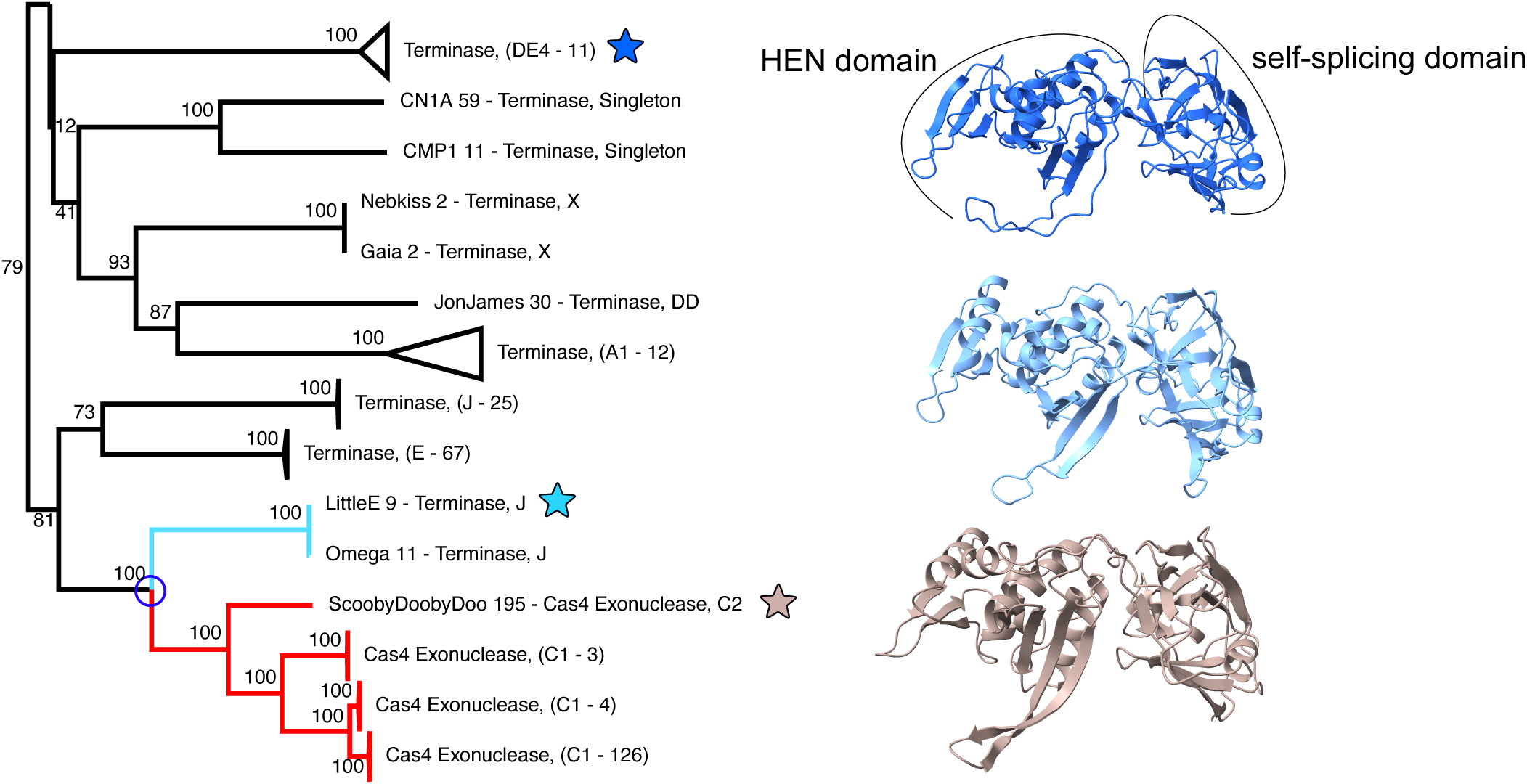
Subtree from phylogeny of class 3 inteins (Figure 4) depicting one instance in which a Cas4 exonuclease intein emerges from a group of terminase large subunit inteins. For each intein indicated with a star (cluster DE4 representative phage APunk terminase large subunit intein, LittleE terminase large subunit intein, and ScobyDoobyDoo (SDD) Cas4 exonuclease intein) a predicted protein structure was generated with AlphaFold. The predicted intein structures (right) are APunk (top) LittleE (middle) and SDD (bottom). Alignments of the predicted intein structures using the Matchmaker tool in ChimeraX produce the following results: LittleE aligned to SDD yields a sequence alignment score of 995.6, and RMSD of 2.716; APunk aligned to LittleE yields a sequence alignment score of 644.9, and RMSD of 3.097; APunk aligned to SDD yields a sequence alignment score of 643.1, and RMSD of 2.994.

**Figure 6.**
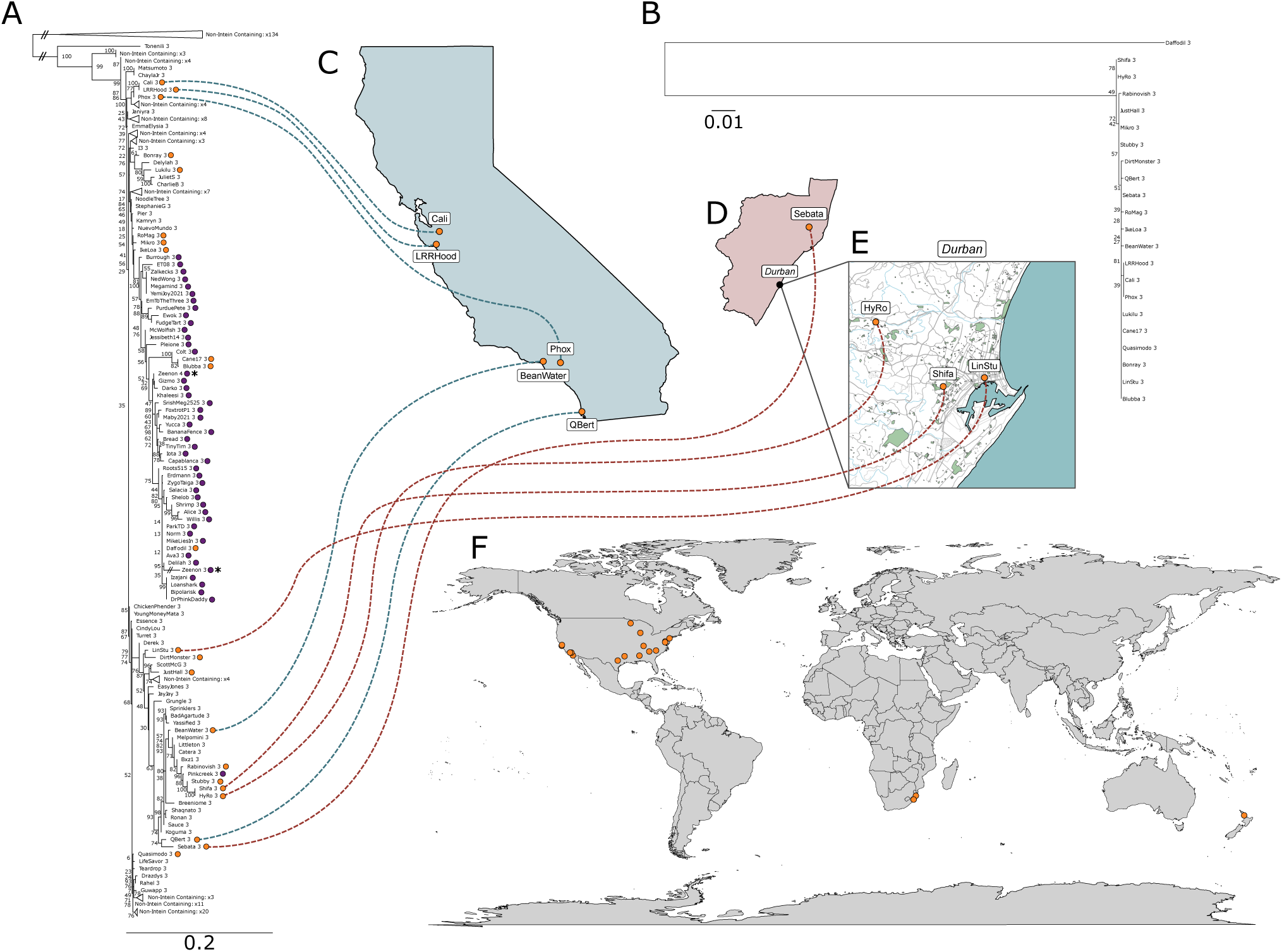
A. Maximum likelihood phylogeny of intein-free and intein-containing nucleotidyltransferase extein-encoding nucleotide sequences, using TIM+F+R5 as the substitution model. Double slashes indicate branches shortened for readability. Circles to the right of taxa names indicate intein presence, with orange and purple denoting two different intein clusters. Ultrafast bootstrap support is listed on internal nodes. There is a split intein in this dataset, from phage Zeenon, indicated by asterisks on the extein phylogeny since its extein is split into two separate open reading frames by the split intein. Scale bar measures substitutions per site. **B**. Maximum likelihood phylogeny of the intein-encoding sequences from the orange cluster, using HKY+F+I as the substitution model. Bootstrap support is shown as it was in **A**. Geographic maps show the distribution of inteins from the orange cluster within: California (**C**), Natal South Africa (**D**), the city of Durban (**E**), and the whole world (**F**). Note that the phages from South Africa (**E**) have identical or nearly identical intein sequences (**B**) but that their exteins do not group together and are separated by exteins harboring no intein, or inteins from a different cluster (**A**). Similarly, the phages isolated in California have nearly identical inteins (**B**), but their exteins are divergent and do not group together (**A**).

### Intein Dissemination within Local Areas

Typically, copies of a particular intein-invaded extein in different individuals will have higher sequence similarity than the inteins that invade them. This is due to the types of genes invaded by inteins often being of vital function to their hosts and thus highly conserved, as well as the increased evolutionary rates experienced by homing elements such as inteins (Goddard and Burt 1999). We further discuss the relative sequence conservation and selection pressures on intein and extein sequences in Supplemental Materials under the section Supplemental Results and Discussion: Sequence Conservation in Inteins and Exteins. Thus, two individuals having less similar exteins but nearly identical inteins is cause for further investigation, as this can indicate a recent transfer event since the intein has not yet had time to diverge from its origin copy despite the higher evolutionary rates of inteins. Geographic data regarding host phage isolation paired with these events can be further investigated to assess the locality of such recent invasion events.

Following this line of analysis, we previously found inteins which appeared to be proliferating into divergent phage extein sequences within local geographic areas (Gosselin et al. 2023). We chose to investigate our dataset for additional cases of local dissemination and proliferation of inteins into new phage populations. We filtered our data in two stages: first to find clusters of closely related inteins (97.5% similarity), then to discover clusters within the filtered set whose extein sequences were more divergent from each other than their respective intein sequences. These clusters were used to construct phylogenetic trees, and their geographic positions were plotted using metadata from PhagesDB. Of the resulting 13 clusters of intein sequences, we focused on three cases which displayed high levels of incongruence between the extein and intein phylogenies during a visual inspection, suggesting recent intein transfer between divergent hosts.

First, we investigated a set of inteins which were found in nucleotidyltransferases (Figure 6). This group of inteins had a global distribution, and of particular interest to us, two groups of co-localized inteins. One group of 5 inteins isolated from California, and a second group found in Durban, South Africa. In both of these locations, inteins exhibited nearly identical nucleotide sequences, but were found in comparatively divergent exteins. Furthermore, these extein sequences were separated in the extein phylogeny by non-intein containing sequences.

We repeated this analysis with a second group of inteins from Ohio and Pennsylvania, found in minor capsid proteins (Figure 7, red cluster). These 7 inteins possess an anti-sense LAGLIDADG HEN. These inteins, despite being identical to each-other (for 5 of the seven), are once again found in divergent extein sequences which were separated from each-other on the phylogenetic tree with non-intein containing taxa, and taxa with completely different inteins.

**Figure 7.**
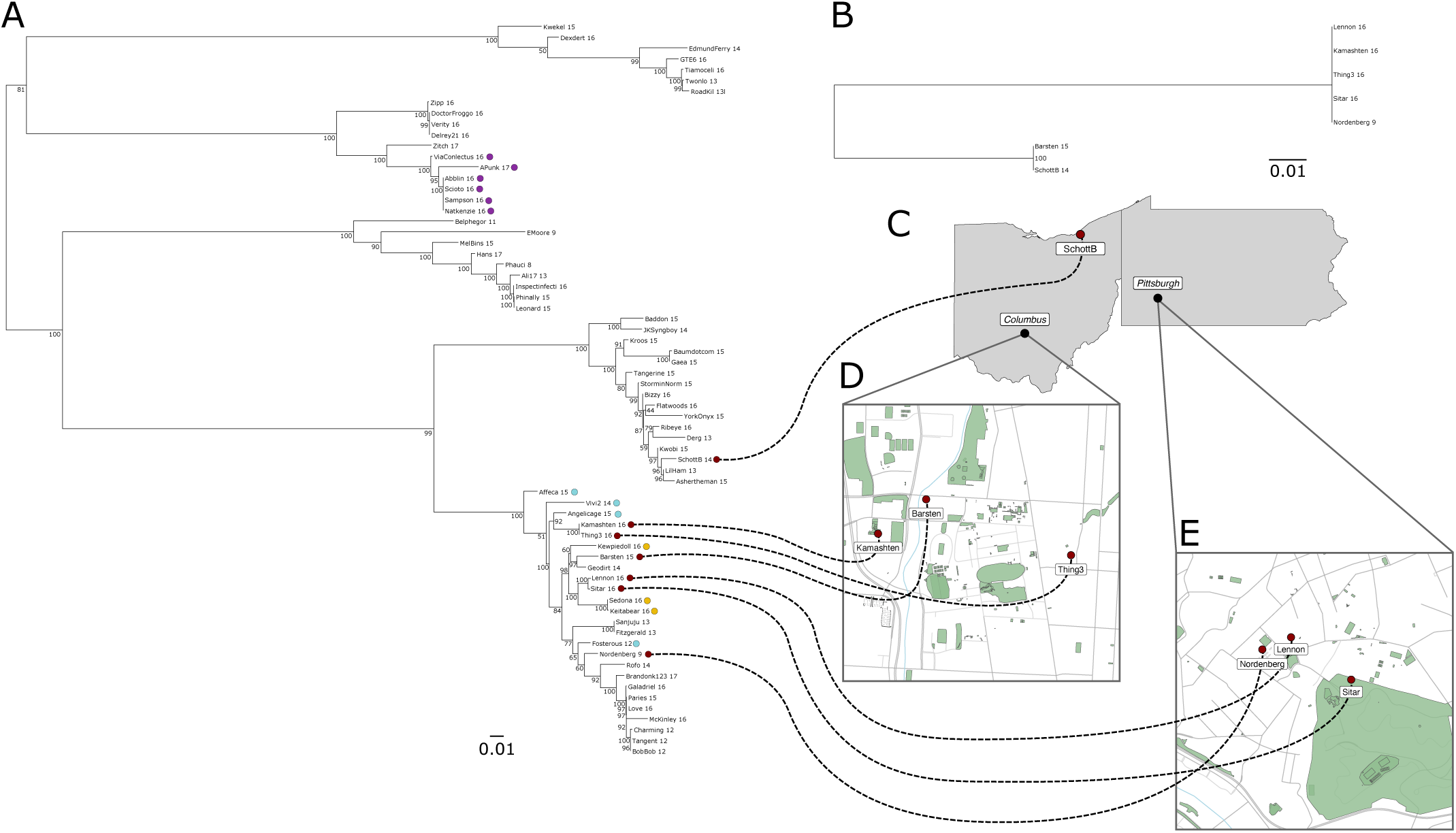
A. Maximum likelihood phylogeny of the nucleotide sequences of intein-containing and intein-free minor capsid proteins using TN+F+R3 as the substitution model. Colored circles to the right of taxa names indicate intein presence, and each color corresponds to a different intein cluster. Ultrafast bootstrap support is listed on internal nodes. Scale bar measures substitutions per site. **B.** Maximum likelihood phylogeny of the intein nucleotide sequences from the red cluster only, using F81+F as the substitution model. Bootstrap support is shown as it was previously. The geographic maps indicate the distribution of inteins from the red cluster in the Ohio-Pennsylvania area (**C**), Ohio State University and surrounding area (**D**), and the University of Pittsburgh and its surrounding area (**E**). Note that the three phages depicted in panel **E** have identical intein sequences, but that the extein from phage Nordenberg diverges from the exteins in Lennon and Sitar and that the three exteins do not form a clan.

Finally, we investigated the inteins found within exteins annotated as putative integrin-like proteins (Figure 8). Once again, these inteins were found in divergent exteins separated on the phylogenetic tree by non-intein containing taxa. Many of these inteins were found specifically within the grounds of Howard University in Washington DC; however, despite the small geographic distance between the phages found there, the exteins showed significant divergence. Furthermore, inteins from each of the three clusters were found on this campus. The vast majority of the inteins from the bottom clan of the intein phylogeny (Figure 8B, gold) were present predominantly on the eastern shoreline of the USA. The clan at the top (Figure 8B, blue) had a much wider distribution — mostly confined to the Mississippi River Valley, and the Midwest. The last of the three clans (Figure 8B, red) was found across the USA.

**Figure 8.**
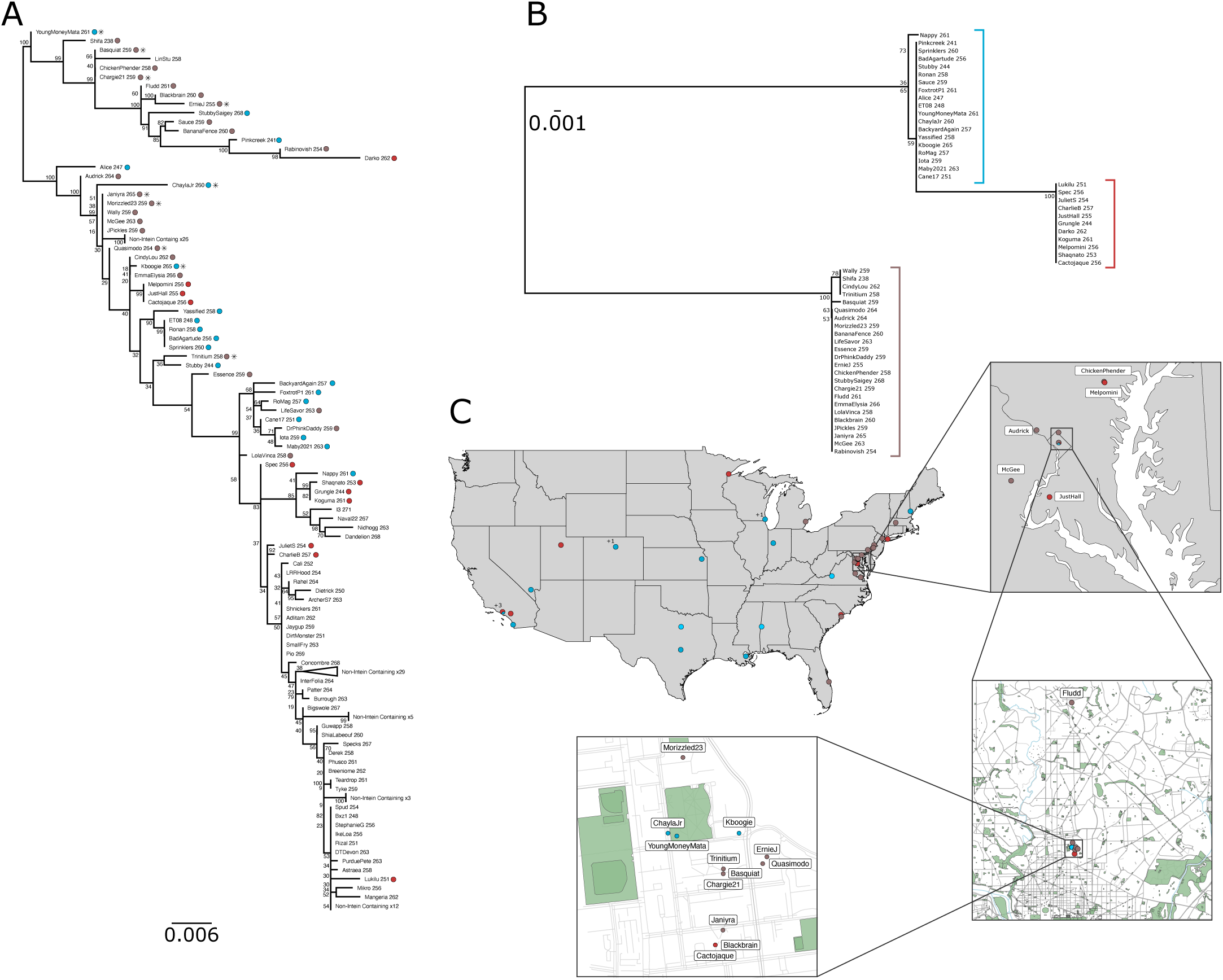
A. Maximum likelihood phylogeny based on the extein-encoding nucleotide sequences of intein-free and intein-containing putative integrin-like proteins using TIM2+F+I+G4 as the substitution model. Colored circles to the right of taxa names indicate intein presence, and the color indicates which cluster (**B**) said intein belongs to. Ultrafast bootstrap support is listed on internal nodes. An asterisk indicates that the phage was isolated within Washington DC. Scale bar measures substitutions per site. **B.** Maximum likelihood phylogeny of all intein-encoding nucleotide sequences from **A**, using TN+F as the substitution model. Colored bars on the three clans correspond to the intein colors in **A**. Bootstrap support is shown as it was previously. **C.** Geographic maps indicating the distribution of these inteins. First within the entire United States of America, and then gradually zooming in on Washington DC, and eventually Howard University. Isolation locations where more than one intein containing phage were collected are indicated with a plus sign and the corresponding number of additional inteins. The color of the circles corresponds to the clans in **B**.

## Discussion

### Overarching Trends

As in previous studies, the inteins we discovered within actinobacteriophages are found predominantly within DNA-binding extein sequences (Kelley et al. 2016). While the majority are within terminase large subunits, the inteins we discovered show a remarkable range of host proteins. In addition to ubiquitous phage genes (such as the terminase large subunit), we find inteins within phage genes with more limited distributions such as thymidylate synthase, or ribonucleotide reductase subunits. Even genes that lack nucleotide-associated enzymatic activity, the minor capsid proteins, were found within the set of invaded phage proteins. Interestingly, the inteins invading minor capsid proteins were exclusively inteins with non-canonical HENs.

Many inteins in our dataset had sporadic distributions across their extein phylogenies, and phylogenies of the intein sequences frequently conflicted with the phylogeny of their extein. We believe that many of these inteins are actively moving between different hosts. However, exceptions do exist in cases of mini inteins (an example of which can be found in Supplemental Figure 8). In addition, inteins that persist in clans of extein sequences, at first sight might be considered to have decaying HENs, and like mini-inteins be vertically inherited (see the purple intein group in Figure 7). However, while the exteins show some sequence divergence, the fact that the inteins are identical likely illustrates a recent invasion event (see below for further discussion).

The many intein-free homologs accompanying intein-containing genes in our dataset indicate that intein homing into new empty target site must be balanced by the loss of inteins in other homologs. Given the seemingly rapid dissemination of inteins into local phage populations, intein loss might also be assumed to occur frequently. This could occur via gradual decay and recombination of the intein, or be the consequence of a fitness cost imparted by the intein on the phage. Inteins are known to cause a drop in fitness; in the archaeon *Haloferax volcanii,* a DNA polymerase intein caused a 7% drop in fitness for the host (Naor et al. 2016). Thus, the alternative loss scenario would act at the population level — the intein imparts enough of a fitness cost such that the intein-containing phage is out-competed by intein-free phages, and the intein containing phages would be slowly selected out of the population. This process would be enhanced if the non-invaded phages evolve at least a partial resistance to intein propagation (perhaps via mutations to the HEN target site). This potential impact on host fitness is a mechanism worth exploring in future work.

### The Nature of Non-Canonical Inteins

In this study we described several inteins with previously unseen, or extremely rare architectures in relation to their HEN domains. To our knowledge, only the anti-sense LAGLIDADG HEN has been reported on in prior literature within the context of a fully functional intein (Gorbalenya 1998; Gogarten et al. 2002). These anti-sense LAGLIDADG HEN containing inteins were discovered in non-phage genomes within the *dnaX* gene of a *Synechocystis* species, and the SpoVR-like protein from *Chloroflexus aurantiacus* J-10-fl (sequences deposited in the InBase database (Perler 2002)). Hence, while not entirely novel, we show that this intein architecture is present within at least one other extein sequence, and importantly, that the intein and anti-sense HEN appear to work together to facilitate ongoing intein invasion events.

VSR-like HEN-containing inteins have been reported on previously, within split-inteins (Dassa et al. 2009). Our findings show that anti-sense VSR-like HEN-containing full inteins also exist and are not merely a point of speculation. If the theory of split-intein evolution outlined by Dassa et al. 2009 is true, then perhaps these newly discovered inteins represent one of the potential off-tracks within their model. It should also be noted here, that Dassa et al. 2009 report that the VSR-like HENs they discovered were related to freestanding HENs within their database (a fact whose importance will be discussed below).

Finally, the endonuclease VII-like HEN containing intein appears to be completely novel, with no known reports. Like the VSR-like HENs, this endonuclease has been reported prior in a free-standing context (Pope et al. 2013).

Given the ubiquity of freestanding HEN genes within actinobacteriophages, the frequency with which these genes seem to proliferate (Edgell et al. 2010; Pope et al. 2013; Zhang et al. 2017; Hatfull 2020; Barth et al. 2023), and the presence of free-standing HENs related to those found within our intein sequences, we believe that the three intein architectures described above originate via invasion of an intein sequence by a free-standing HEN. Given the seeming rarity of these elements, we further propose (in line with the model laid out by Dassa et al. 2009 (Dassa et al. 2009)), that successful integration of these HENs into the region of an intein between splicing associated blocks is extremely rare. The breaking of, and subsequent integration into the intein CDS by a free-standing HEN in this fashion may be relatively common, but most of the time this integration is likely to result in split-inteins, or introduce nonsense mutations which prevent post-translational rescue of extein function via intein splicing.

Assuming the free-standing HEN’s integration into the intein CDS is successful, the path forward is unclear. It is possible the HEN becomes a pseudo-gene, but there have been prior examples of phage genes with nested HEN open reading frames (ORFs) that still translate both proteins (Gibb and Edgell 2007; Wright et al. 2021). In our cases, the secondary ORFs typically lie outside the bounds of the intein splicing domain, and given the splicing domain’s high conservation, it is probable that the spicing domain still functions properly. Additionally, many of these non-canonical inteins were found in the crucial terminase large subunit gene, and it is therefore unlikely that the intein is not splicing itself out — these phages would otherwise be unable to properly package their DNA into their capsid (a certain end for that phage lineage). Our phylogenetic reconstructions of the anti-sense LAGLIDADG-containing inteins provided evidence for continued invasion. These inteins are present in multiple locations within divergent host exteins, and importantly are separated with meaningful support by relatives that lack the intein. Such a pattern is not reasonably explained via vertical inheritance and loss, and it is therefore likely that the splicing domain and anti-sense HEN domain of that particular intein still function as a single unit. This may also be true of two groups of inteins with VSR-like HENs (Supplemental Table 1, clusters 60 and 91) which likewise show a pattern of inheritance more in-line with horizontal gene transfer. However, the other cases of VSR-like HEN-containing inteins exhibit patterns suggestive of vertical inheritance (or were singletons and thus no evolutionary inference could be drawn). The endonuclease VII-like HEN-containing inteins presented a more ambiguous distribution. While most are clustered together, there are a number of closely related extein alleles without an intein. This, plus the sequence similarity of these inteins (for example, the inteins of cluster 47 — which includes phages Angelicase, Affeca, Keitabear, and Sedona — are all identical with the exception of Angelicase), suggests that these inteins may still be transferring between phages horizontally (Supplemental Figure 9, Supplemental Figure 4).

If these inteins can still splice, but can no longer successfully jump, then selection will likely lead to these secondary ORF’s very quickly accumulating stop codons and ceasing to function. This would be especially likely in the case of anti-sense ORFs since they would not benefit from the purifying selection acting upon the host extein to remove nonsense mutations. Mini-inteins and split-inteins would be the most likely observable outcome of this process. While we do find nine inteins which appear to have decaying HEN domains, and a further nine mini-inteins, a survey strategy fine-tuned towards finding split and mini inteins would be needed to look for these potential relics of decay. Potentially, the secondary ORFs encoded on the same strand could transit into the extein’s reading frame via one or two base pair indels, resulting in a fully contiguous intein with a novel HEN.

Inteins created by the mechanism described above thereby provide meaningful support for a “protein splicing first” theory of intein evolution (Derbyshire and Belfort 1998; Gogarten et al. 2002; Dassa et al. 2009; Green et al. 2018). The number of intein-associated HEN’s (including HNH, VSR-like, GIY-YIG, LAGLIDADG, etc. (Dalgaard, Klar, et al. 1997; Gorbalenya 1998; Gimble 2000; Dassa et al. 2009)) suggests that there have been multiple separate invasions of intein coding sequences by free-standing HENs (Derbyshire and Belfort 1998). Perhaps most, if not all, of these HENs came into intein sequences via a free-standing HEN, proliferated through the intein’s self-splicing machinery, and were shifted by an indel or an inversion into a contiguous orientation.

### Frequent Horizontal Transfer of Inteins

Extein phylogenies with intein presence mapped onto them, such as in Figures 6A, 7A, and 8A, can reveal likely cases of vertical inheritance, recent dissemination, and horizontal transfer of inteins.

Using an extein phylogeny with intein presence mapped will differentiate between horizontal transfer and vertical inheritance or recent dissemination; however, intein phylogenies and assessment of intein and extein conservation are required to differentiate between vertical inheritance with frequent intein loss events and recent dissemination.

A set of intein-containing exteins which all group together initially indicates vertical inheritance of that intein, such as the group in Figure 7A containing phage ViaConlectus. While the phylogeny is unrooted, if one follows the presented arrangement one would infer that ViaConlectus was invaded by the purple cluster intein, and this intein was then vertically inherited by phages APunk, Abblin, Scioto, Sampson, and Natkenzie. To assess whether the event is a case of vertical inheritance or recent rapid dissemination, the degree of similarity between inteins in these cases must be further analyzed, as well as geographic proximity between hosts when such information is available. In such cases, extremely similar inteins in geographically-close but less similar exteins may indicate a recent intein dissemination event.

In contrast to the phylogenetic patterns suggesting vertical inheritance or recent local dissemination are cases where we find inteins within extein sequences that lack a direct common ancestor and are separated on the extein phylogeny by many nodes leading to other non intein-containing extein sequences. In such cases, a vertical inheritance pattern would require many loss events to be an accurate model given these phylogenies. To supplement findings from extein phylogenies with intein presence mapping, the phylogeny of the intein of interest can be reconstructed and its topology compared to that of the extein phylogeny to identify incongruencies, which often provide further evidence of a non-vertical inheritance pattern. For example, in the extein phylogeny with intein presence mapping in Figure 7A, we see a well-supported (99% ultrafast bootstrap support) split between a group of exteins from phages including phage SchottB, and a group of exteins from phages including phages Kamashten, Thing3, Barsten, Lennon, Sitar, and Nordenberg. Despite the evolutionary distance between exteins, these phages all contain near identical inteins at the same position in their extein (inteins from the red cluster in Figure 7A). To supplement this observation, Figure 7B depicts the intein phylogeny for this red cluster of inteins which shows the SchottB intein grouping closely with the Barsten intein, despite the large well-supported evolutionary distance between their exteins. The most parsimonious model explaining this observation is a horizontal transfer event between SchottB and a member of the Kamashten group, as opposed to vertical inheritance with many independent loss events. Applying this phylogenetic analysis to each of our extein-intein phylogeny pairings thus reveals a multitude of cases for the horizontal transfer of inteins between phages.

For highly similar inteins which possess a wide geographic distribution, the next question inevitably becomes how did they get there? While humans are almost certainly acting as a secondary or tertiary vector for the intein, we propose that these inteins are likely dispersing in part via the phages we find them in and in part, via the phage’s host bacteria. Previous literature, and our own anecdotal observations, find that many phage-associated inteins are found inside of bacterial genomes, most likely within prophages (25). Thus, inteins experience increased genetic mobility, taking advantage of both the viral and bacterial vectors. As a prophage, the intein can propagate not only when the bacterial cell replicates, but also through the numerous avenues of horizontal gene transfer that bacterial cells engage in (Soucy et al. 2015), and by transfer into closely related phages upon infection.

### Geographic Spread of Inteins

Our dataset contains several groups of inteins which are extremely similar on the nucleotide level (and in the cases described, often identical), these inteins must have diverged from one another relatively recently. This presents two options: rapid host proliferation with direct vertical inheritance, or rapid intein proliferation via horizontal gene transfer. For some of the inteins in our dataset, the former may be considered a possibility (see the clan of intein containing exteins colored purple in Figure 7); however, where the intein containing exteins form a well-supported clan with limited geographic distribution, the observation that the inteins are identical, whereas the exteins are divergent, suggests that the inteins have recently invaded the divergent exteins instead of being vertically inherited. This argument is strengthened by the observation that inteins found in divergent exteins experience less selection for sequence conservation than the exteins that harbor them (see Supplemental Materials: Supplemental Results and Discussion – Sequence Conservation in Inteins and Exteins). The finding of nearly identical inteins found in divergent exteins in close geographical proximity suggests that intein invasion represents an example of a highly effective gene drive acting on the local level following the introduction of a new founder intein allele. The forces that limit this drive (after all, not all homologous exteins within a given location contain the intein) need further study (see the above discussion of fitness cost to the host that can be associated with inteins). Phages, because of their rapid propagation and large population sizes provide an excellent opportunity to study intein population dynamics in more detail.

### Nucleotide binding sites as an avenue for extein-switching

The instance of Cas4 exonuclease inteins evolving from terminase large subunit inteins on the class 3 intein phylogeny involving phages LittleE and ScoobyDoobyDoo (SDD) (Figure 4) provides a compelling argument for the use of highly similar nucleotide binding sites as a means of intein movement between otherwise dissimilar genes. The stretch of identical nucleotides between the two intein insertion sites (10 bases on the 5’ side, 2 bases on the 3’ side) in conjunction with the other shared positions within the +/-30bp surrounding the site (Supplemental Figure 7A) may be sufficient for recognition and invasion by the same intein. The proposed range for HEN recognition sites is ∼12-40bp, with recognition sites in smaller genomes such as phage genomes being on the lower end of the range (Jurica and Stoddard 1999). Additionally, many types of phage recombinases can tolerate some dissimilarity between regions used for homologous recombination (Lopes et al. 2010) suggesting this intein could possibly invade either site if given the chance.

To further complete the picture and investigate why the SDD terminase large subunit and LittleE Cas4/RecB-like exonuclease are intein-free, we analyzed the regions corresponding to the intein-insertion sites on the intein-free proteins (Supplemental Figure 7B) and find SDD’s Cas4 exonuclease gene to be a more hospitable landing pad for the LittleE terminase large subunit intein by metrics of insertion site recognition and residues at the insertion site required for intein self-splicing (the SDD terminase large subunit encodes an alanine at the position which would need to be a serine, threonine, or cysteine for successful intein splicing to occur) (Supplemental Figure 7B) (Mills et al. 2014; Turgeman-Grott et al. 2023). Additionally, LittleE is a temperate phage of the same strain of bacteria that SDD is a lytic phage of (*Mycobacterium smegmatis* mc2155) (Russell and Hatfull 2017).

The presented evidence taken together suggest a model in which the recipient genome was exposed to the donor’s terminase large subunit intein during co-infection or infection of a host containing the donor’s prophage. Then, either homing of the donor terminase large subunit intein into the recipient’s terminase large subunit gene was thwarted by the lack of similarity between these terminase large subunits, or homing was successful but nonviable due to lack of a splicing-compliant residue on the C-terminal side of the insertion site. By happenstance, a region of similarity to the donor terminase large subunit present in the recipient Cas4 exonuclease gene conferred by a shared nucleotide binding motif provided a viable alternate insertion site for the intein (Figure 9). Being in a brand-new context also lets this copy of the intein begin a fresh evolutionary trajectory, as the majority of homologous genes in the population will initially lack this intein. This may explain the presence of SDD’s Cas4 exonuclease intein in over 130 other phage Cas4 exonucleases (Figure 4). As PhagesDB and other phage genome sequence resources continue to grow, increased sampling will tell a clearer story of the events that unfolded in the evolution of these inteins.

**Figure 9:**
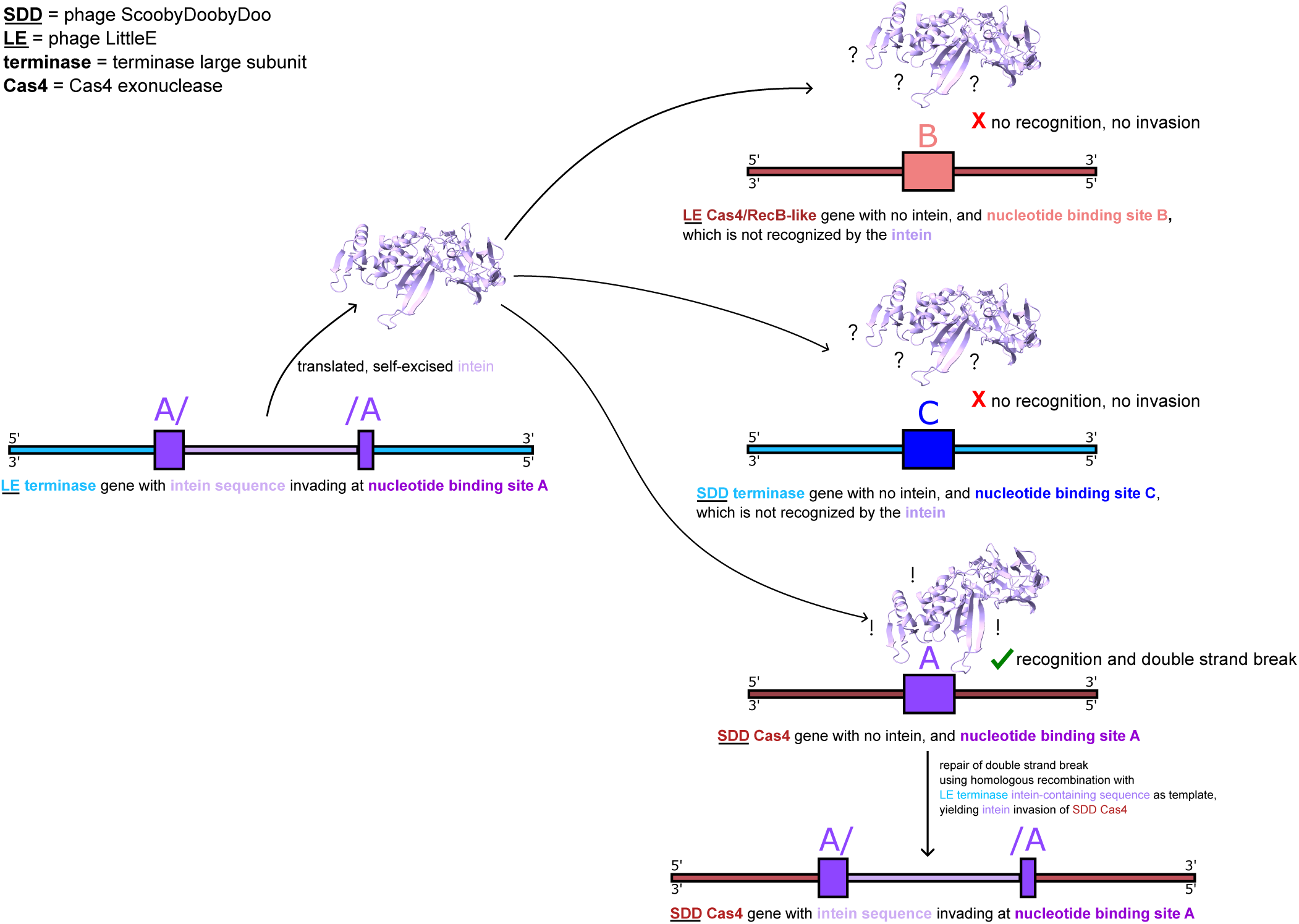
Three different scenarios for attempted intein invasion by the LittleE terminase large subunit (TLS) intein, explaining simultaneous presence of intein in LittleE TLS and ScoobyDoobyDoo (SDD) Cas4 exonuclease genes, but lack of intein in LittleE Cas4/RecB-like endonuclease and SDD TLS genes. TLS exteins are depicted as thin blue boxes, Cas4 exonuclease exteins are depicted as thin red boxes, and the intein sequence within invaded genes is depicted as a thin purple box. Nucleotide binding site A (depicted as a thick purple box) is the insertion site recognized by the LittleE TLS intein, and is present in both the LittleE TLS gene and the ScoobyDoobyDoo (SDD) Cas4 exonuclease gene. Nucleotide binding site B (depicted as a thick light red box) is not recognized by the LittleE TLS intein, and is present in the LittleE Cas4/RecB-like exonuclease gene in place of nucleotide binding site A. Thus, the LittleE TLS intein cannot engage in homing with the LittleE Cas4/RecB-like exonuclease gene and invasion cannot occur. Nucleotide binding site C (depicted as a thick dark blue box) is not recognized by the LittleE TLS intein, and is present in the SDD TLS gene in place of nucleotide binding site A. Thus, the LittleE TLS intein cannot engage in homing with the SDD’s TLS gene and invasion cannot occur. Since the SDD Cas4 exonuclease gene contains nucleotide binding site A, the LittleE TLS intein can engage in homing, allowing the invasion of the LittleE TLS intein into the SDD Cas4 exonuclease gene.

## Conclusions

The work outlined above shines a light on the previously unknown breadth of inteins within Actinobacteriophage genes. Throughout this research we repeatedly find disjunct distributions of intein containing exteins in extein phylogenies, and propose that the conflict between extein and intein phylogenies suggests that inteins within Actinobacteriophages frequently home into target sites within intein-free genes. We find evidence for multiple cases of rapid geographically local episodes of intein invasion that support the prior assertion. In addition, we describe several previously unseen or rare intein architectures, and provide a hypothesis to explain their potential occurrence and apparent continued dissemination. We also describe a model for how inteins may home into new target sites, within previously un-invaded gene families, based on the intein phylogenies and the similarities and dissimilarities between target sites.

Finally, we wish to stress that these non-canonical inteins may be significantly more common than even what we have outlined here. Given their overlapping ORFs, these inteins are likely missed in automated annotation pipelines, especially within phages — current SEA-PHAGES training argues for “…typically only one frame in one strand is used for a protein-coding gene…” (SEA PHAGES). Even to the trained eye, these inteins may currently be glossed over as ‘decaying’ canonical inteins or misclassified as mini-inteins. Significant work will be needed to confirm that inteins listed as decaying are correctly classified, and to discover the true extent of these inteins within phages and their bacterial hosts.

## Materials and Methods

### Code Availability

All code used in this project is available at https://github.com/sophiagosselin/intein_distribution along with usage information. The two exceptions to this are ICE-BLAST and Invader-Sim which are kept in separate repositories detailed below.

### Iterative Cluster Expansion BLAST: ICE-BLAST

To increase efficiency in mass intein sequence retrieval, we developed “Iterative Cluster Expansion BLAST” (ICE-BLAST). ICE-BLAST is a Perl (v 5.36.0) based program using PSI-BLAST (Altschul et al. 1997) as the principal sequence search tool and USEARCH clustering (Edgar 2010) to identify new queries from previous iterations. Through this process, ICE-BLAST retrieves highly divergent homologous protein sequence matches. In short, ICE-BLAST iteratively searches amino acid databases with PSI-BLAST, clusters filtered outputs using USEARCH, then uses the representative sequence of each match cluster (centroids) as the next query sequences (Figure 10A).

**Figure 10.**
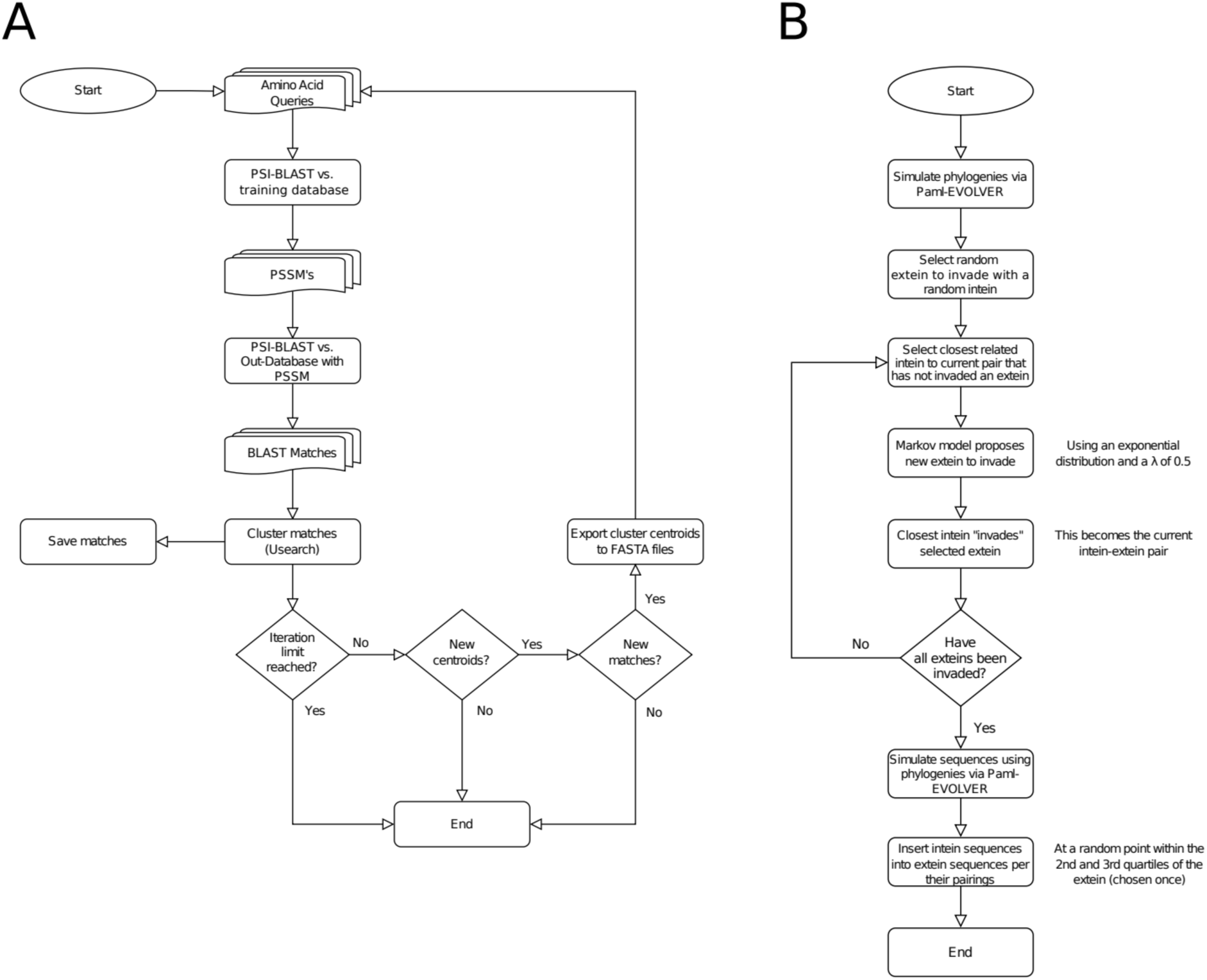
Flow charts describing the algorithms of ICE-BLAST and Invader-Sim. **A.** ICE-BLAST begins by taking amino acid query sequences and iteratively searching against a training amino acid sequence databank using PSI-BLAST. The matches resulting from this iterative search are clustered by sequence similarity, where a representative amino acid sequence is selected from each cluster (referred to as the centroid sequence for that cluster). The centroids of each cluster are then used as input for a new round of PSI-BLAST searches, with the process repeated until either the iteration limit is reached, there are no new centroids, or there are new centroids but no new matches over all. **B.** Invader-Sim begins by using PAML-EVOLVER to simulate an extein phylogeny and an intein phylogeny. From the phylogeny of potential exteins, a random individual is chosen to be invaded by an intein from the intein phylogeny. The closest relative of the randomly selected intein is given a new extein to invade, with the extein selection proposed by a Markov model. The process is repeated until all exteins are invaded by an intein. Simulated sequences are then generated using PAML-EVOLVER.

ICE-BLAST is available at https://github.com/sophiagosselin/ICE_BLAST along with detailed usage, dependencies, and other information. Algorithm details and parameters used for this study are available in Supplemental Materials.

### Simulations of Intein Evolution: Invader-Sim

Since ICE-BLAST involves multiple individual searches, one must take proper precautions against false-positives. False-positives can introduce errors that will propagate through all downstream iterations. Hence, it was important to test ICE-BLAST against known data to assess overall sensitivity and specificity. We developed a Perl (v 5.36.0) based sequence simulation program, Invader-Sim, to generate sequences simulating intein invasion into extein sequences. The program uses a combination of PAML-EVOLVER (v4.9j) (Yang 2007) and a Markov chain to simulate datasets of intein-containing extein sequences (Figure 10B). ICE-BLAST performed as well as PSI-BLAST on the simulated datasets. Results and additional discussion can be found in Supplementary Materials.

Invader-Sim is available at https://github.com/sophiagosselin/invader_sim. Information on the method and parameters used for this study are available in the Supplemental Materials.

### Dataset Curation: Applying ICE-BLAST to PhagesDB

Using PhagesDB data (Russell and Hatfull 2017), two databases were constructed: one from amino acid sequence files, and one from whole genomes, with associated metadata from the PhagesDB API. The genomic and amino acid databases were downloaded and formatted on September 28 2023 and November 2 2023 respectively. The databases were not updated over the course of this work, due to reverse incompatibility of PhagesDB updates.

The initial ICE-BLAST query set for this study consisted of 78 intein sequences drawn from three sources: InBase (final update 2010) (Perler 2002), known Actinobacteriophage inteins (Kelley et al. 2016), and inteins found by the Gogarten Lab in the PhagesDB database between 2020 and 2024 (Gosselin et al. 2023) (Supplemental Table 3). Initial PSSMs were trained on Uniref50 (Suzek et al. 2015) downloaded November 7 2021. For the ICE-BLAST search, a percent identity of 60%, an e-value of 1e-20, and a domain specific cutoff bound threshold of 25% were used as inputs for filtering.

Initial results from ICE-BLAST contained 1994 sequences which were then clustered by USEARCH (v11.0.667) (Edgar 2010) at 70% identity using the cluster_fast algorithm. Associated intein-invaded extein sequences were then extracted from the BLAST database using blastdbcmd. Putative intein sequences were aligned to the invaded extein sequences using MAFFT (v7.490) (Katoh and Standley 2013). When intein identity was uncertain, BLAST against NCBI’s non-redundant database (nrdb) (Altschul et al. 1990), and HHPred against PDB, Pfam, and NCBI CDD (Zimmermann et al. 2018), were used to determine whether the sequence was actually an intein, and where its boundaries laid.

False positives were removed from the dataset at this point. This screening reduced the dataset to 784 confirmed intein-containing sequences (Supplemental Table 1). The large was attributed entirely to a false positive match to a DNA Primase sequence; causing over 1000 matches to DNA primase extein domains which were falsely incorporated. Extein sequences were re-annotated using BLAST and HHPred as described above. Inteins were classified as class 1, 2, or 3, by comparing to self-splicing domain consensus sequences (Tori et al. 2010). In-house Perl scripts were used to extract associated coding sequences from the mentioned PhagesDB genome database. Intein and extein sequence files are available in Supplemental Data.

HENs were classified using the method above, with two exceptions. First, we used the PHROG (Terzian et al. 2021) database within HHPred (Zimmermann et al. 2018) (especially for more divergent sequences) to identify potentially related HENs as specified in the PFAM section of its entries. Second, we used InterProScan (as of August 2024) (Jones et al. 2014) when BLAST and HHPred were inconclusive.

### Geographic Maps

First, intein nucleotide sequences were clustered using USEARCH (97.5% sequence identity) resulting in 91 clusters. Clusters with less than four members were excluded, as a meaningful phylogenetic tree cannot be generated. If exteins within a cluster were more self-similar than their intein sequences were, the cluster was removed. Remaining clusters were then used to build intein nucleotide phylogenies, and their extein sequences (with intein-less homologs from PhagesDB) were used to construct an extein phylogeny. Only clusters whose intein and extein phylogenies were in-congruent were selected for further study.

Extein phylogenies were marked for intein presence, and the inteins present were then mapped onto their geographic isolation location. Phage isolation locations were extracted from PhagesDB’s API. Maps of intein isolation locations were created using R (v4.4.1) and the following libraries: sf (v1.0-16) (Pebesma 2018), ggplot2 (v3.5.1) (Wickham 2016), maps (v3.4.2) (Deckmyn 2023), mapdata (v2.3.1) (Brownrigg 2022), mapproj (v1.2.11) (Minka and Bivand 2023), rnaturalearthdata (1.0.0) (South et al. 2024a), and rnaturalearthhires (v1.0.0.9000) (South et al. 2024b).

### Phylogenetic Trees and AU Tests

Phylogenies in this study were built from MAFFT sequence alignments. Unless specified, MAFFT was allowed to select the best alignment algorithm for a given set of sequences (strategy: 1. --auto). IQ-TREE (v2.0.7) using ModelFinder (-m MFP) and 1000 ultrafast bootstraps (-bb 1000) (Minh et al. 2020) was used for phylogenetic reconstruction. Models selected are included in phylogeny figure legends.

Trees were visualized using Dendroscope (v3.8.10) (Huson et al. 2007), Figtree (v1.4.4) (Rambaut 2012), and Inkscape (v1.3.2) (Developers). Clans with extremely similar or identical sequences were collapsed in some phylogenies for readability. Un-collapsed versions are included in the supplemental material (see Figure 4 and Supplemental Figure 10 for examples of collapsed and un-collapsed respectively).

AU tests (Shimodaira 2002) were performed via the method built into IQ-TREE2 (Minh et al. 2020).

### Predicted Protein Structures

Protein structures were predicted for non-canonical inteins and their associated secondary ORFs using AlphaFold (Jumper et al. 2021) through UCSF ChimeraX (Pettersen et al. 2021; Meng et al. 2023). Structures were colored according to the predicted local distance difference test (pLDDT) value per-residue. Each predicted structure was used as query in a DALI search of the PDB database (Holm et al. 2023).

AlphaFold (Jumper et al. 2021) was also used to generate predicted structures in the extein switching study. UCSF ChimeraX (Pettersen et al. 2021; Meng et al. 2023) was used to compare the predicted structure of the LittleE terminase large subunit to the solved structure of a phage terminase large subunit bound to ATP (Zhao et al. 2013).

### Using Predicted Structures to Investigate and Classify Non-Canonical HENs

For the inteins with non-canonical HENs, predicted structures suggest that the portion of the intein which should be an in-frame HEN does not make any recognizable structures (Supplemental Figure 1), and cannot be matched to any known structure using the DALI server PDB search tool. However, predicted structures of the anti-sense LAGLIDADG HEN, anti-sense VSR-like HEN, and out-of-frame endonuclease VII-like HEN, all matched to their associated predicted domains (Supplemental Figure 2). It is important to note that these matches were only to the core catalytic domain of these HENs; significant portions of these structures could not be matched to known proteins. For the anti-sense LAGLIDADG HEN, the LAGLIDADG core sits between residues 40 and 185, matching to PDB structure 2O7M (Prieto et al. 2007) (CreI — a known LAGLIDADG HEN) with a Z-score of 14.8. The VSR-like HEN predicted structure matched to PDB structure 1CW0 (Tsutakawa et al. 1999) (VSR) with a Z-score of 10.6 across the residues 144 to 267. Finally, the endonuclease VII-like HEN matches to PDB structure 1E7D (Raaijmakers et al. 2001) (a recombination endonuclease VII) with a Z-score of 6.5 across residue 1 to residue 67.

### Intein Insertion Site Consensus Sequence Plots

WebLogo (Crooks et al. 2004) was used to generate consensus sequence plots in this study.

## Acknowledgments

We would like to thank the students and faculty of the SEA-PHAGES program and community for their work isolating, sequencing, and annotating actinobacteriophages, for maintaining PhagesDB, and for helpful discussions. We would like to thank Dr. Jonathan Klassen, Dr. Daniel Gage, and Dr. Carolyn Teschke at the University of Connecticut for helpful comments, and insights into the non-canonical inteins. Additional thanks to the Computational Biology Core, Institute for Systems Genomics, University of Connecticut, for the use of computational resources.

This work was supported through grants from the National Science Foundation within the BSF-NSF joint research program, NSF/MCB 1716046, and UConn’s OVPR Research Excellence Program (REP-0000000833). The funders had no role in study design, data collection and analysis, decision to publish, or the writing of this manuscript.

