## Supplemental Materials for "Actinobacteriophage Inteins: Host Diversity, Local Dissemination, and Non-Canonical Architecture"

### Supporting Information Note:

The following supplemental materials include extended details regarding our methodology, and some portions of the analysis which were cut for space, but may be of interest to some readers.

### Supplemental Material Table of Contents:


Supplemental Figures 1 – 13 ------------------------------------------------------ pages 3-19

Supplemental Tables 1 – 6 --------------------------------------------------------- pages 20-25

Supplemental Data Description ---------------------------------------------------- page 26

- Identified Inteins - Extein Nucleotide Sequences.zip
- Identified Inteins - Extein With Intein Nucleotide Sequences.zip
- Identified Inteins - Intein Nucleotide Sequences.zip
- Data for Supplemental Figure 3 – Minor Capsid VSR Inteins Aligned.fna
- Data for Supplemental Figure 4 – Endonuclease VII Inteins Aligned.fna
- Data for Supplemental Figure 5 – HNH Inteins Aligned.fna
- Data for Supplemental Figure 6 – Anti-Sense LAGLIDADG Inteins Aligned.fna
- Supplemental Invader Sim Results Graphs.zip

Supplemental Methods Information ------------------------------------------------ pages 27-29

- ICE-BLAST: Detailed Methodology
- Invader-Sim: Detailed Methodology
- Invader-Sim: Study Parameters

Supplemental Results and Discussion -------------------------------------------- pages 30-35

- ICE-BLAST Testing using Simulated Data
- Intein Insertion Site Analysis
- Sequence Conservation in Inteins and Exteins

Supplemental Material References ----------------------------------------------- page 36

### Supplemental Figures

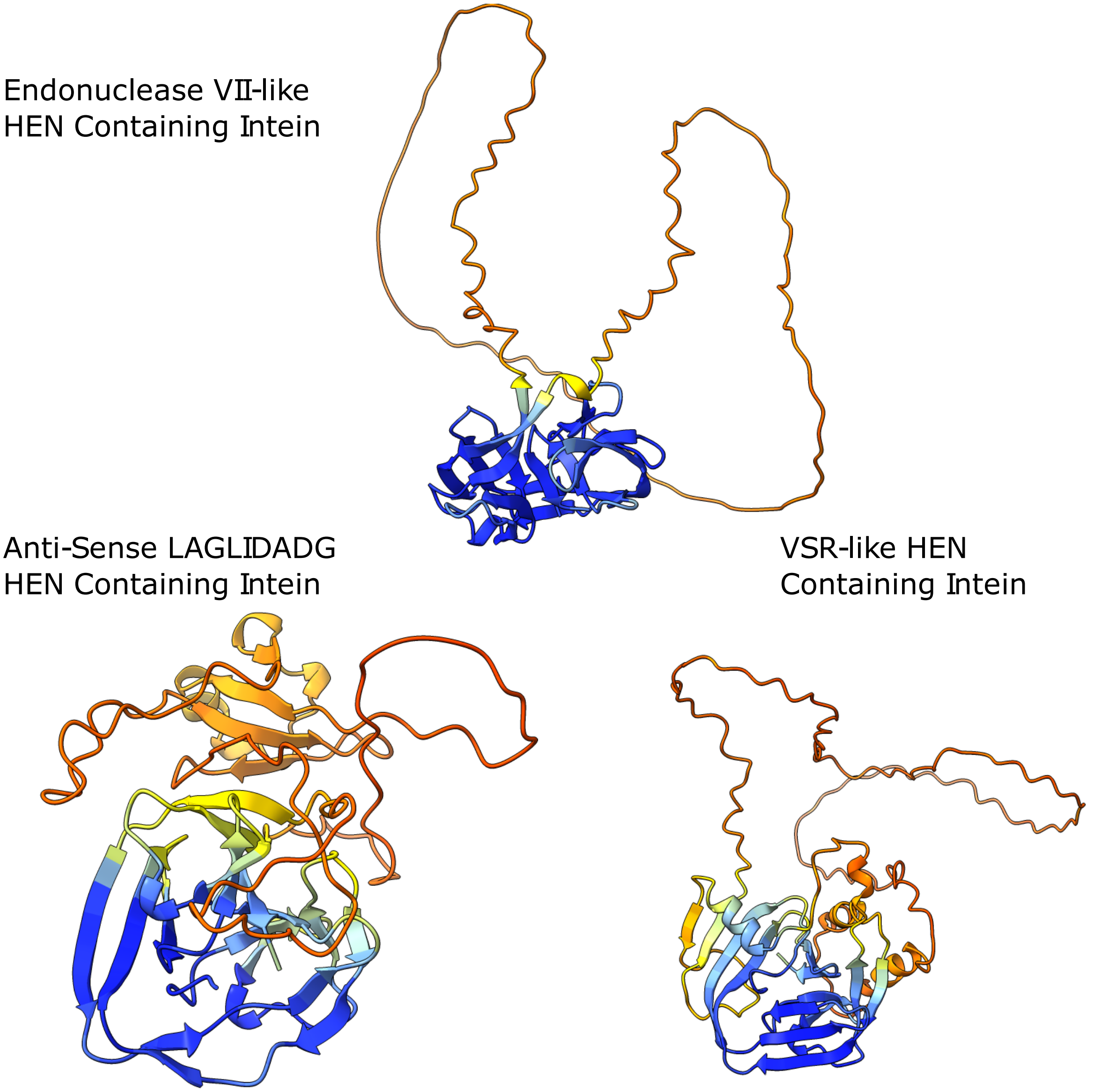


**Supplemental Figure 1:** AlphaFold predicted protein structures of the inteins associated with the VSR-like HEN (from phage Angelicage), the Enodnuclease-VII-like HEN (from phage Affeca), and anti-sense LAGLIDADG HEN (from phage Nordenberg). Predicted structures are colored per site according to predicted local distance difference test (pLDDT) scores. Colors represent: pLDDT > 90 (dark blue), 90 > pLDDT > 70 (light blue), 70 > pLDDT > 50 (yellow), and plDDT < 50 (orange to red). All intein nucleotide sequences are available in Supplementary Data in the Identified Inteins – Intein Nucleotide Sequences section.

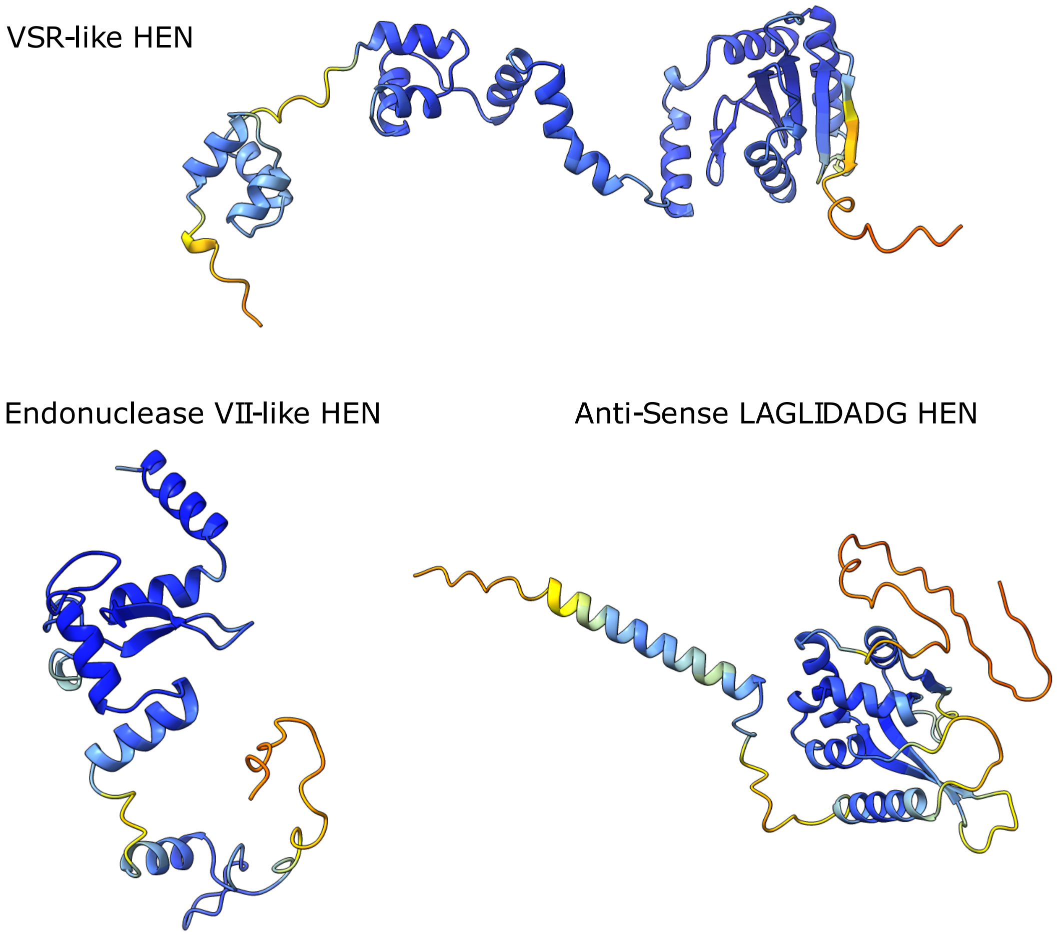


**Supplemental Figure 2**: AlphaFold predicted protein structures of the VSR-like (from phage Angelicase), Enodnuclease-VII-like (from phage Affeca), and anti-sense LAGLIDADG (from phage Nordenberg) HEN's. Predicted structures are colored per site according to predicted local distance difference test (pLDDT) scores. Colors represent: pLDDT > 90 (dark blue),  90 > pLDDT > 70 (light blue), 70 > pLDDT > 50 (yellow), and plDDT < 50 (orange to red). All intein nucleotide sequences are available in Supplementary Data in the Identified Inteins – Intein Nucleotide Sequences section.

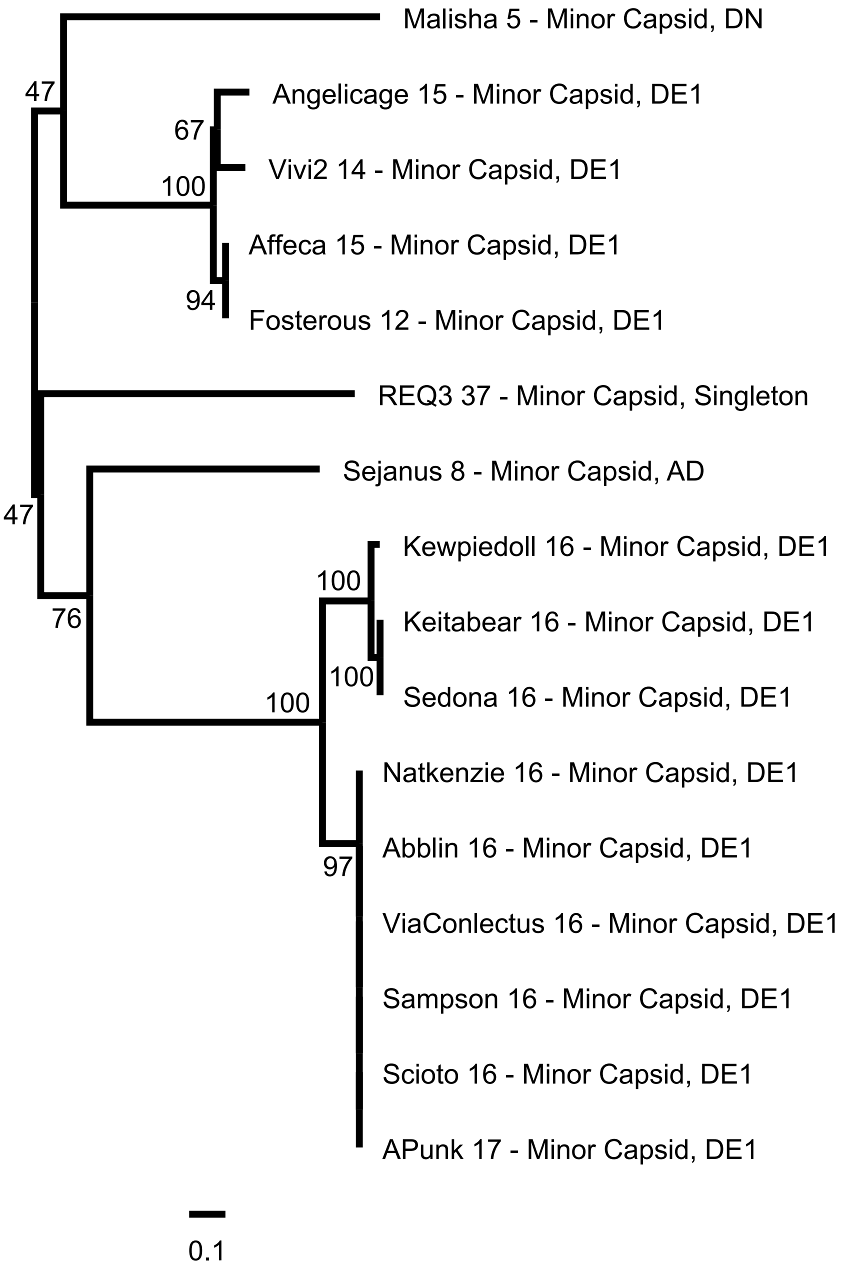


**Supplemental Figure 3**: Maximum likelihood phylogeny of nucleotide sequences encoding inteins with VSR-like HEN's using HKY+F+G4 as the substitution model. Ultrafast bootstrap support is listed on internal nodes. Tip labels include the phage name, gene number, associated extein annotation, and phage cluster. Scale bar measures substitutions per site. Nucleotide sequence alignment is available in the Supplemental Data section under Data for Supplemental Figure 3 – VSR Inteins Aligned.fna

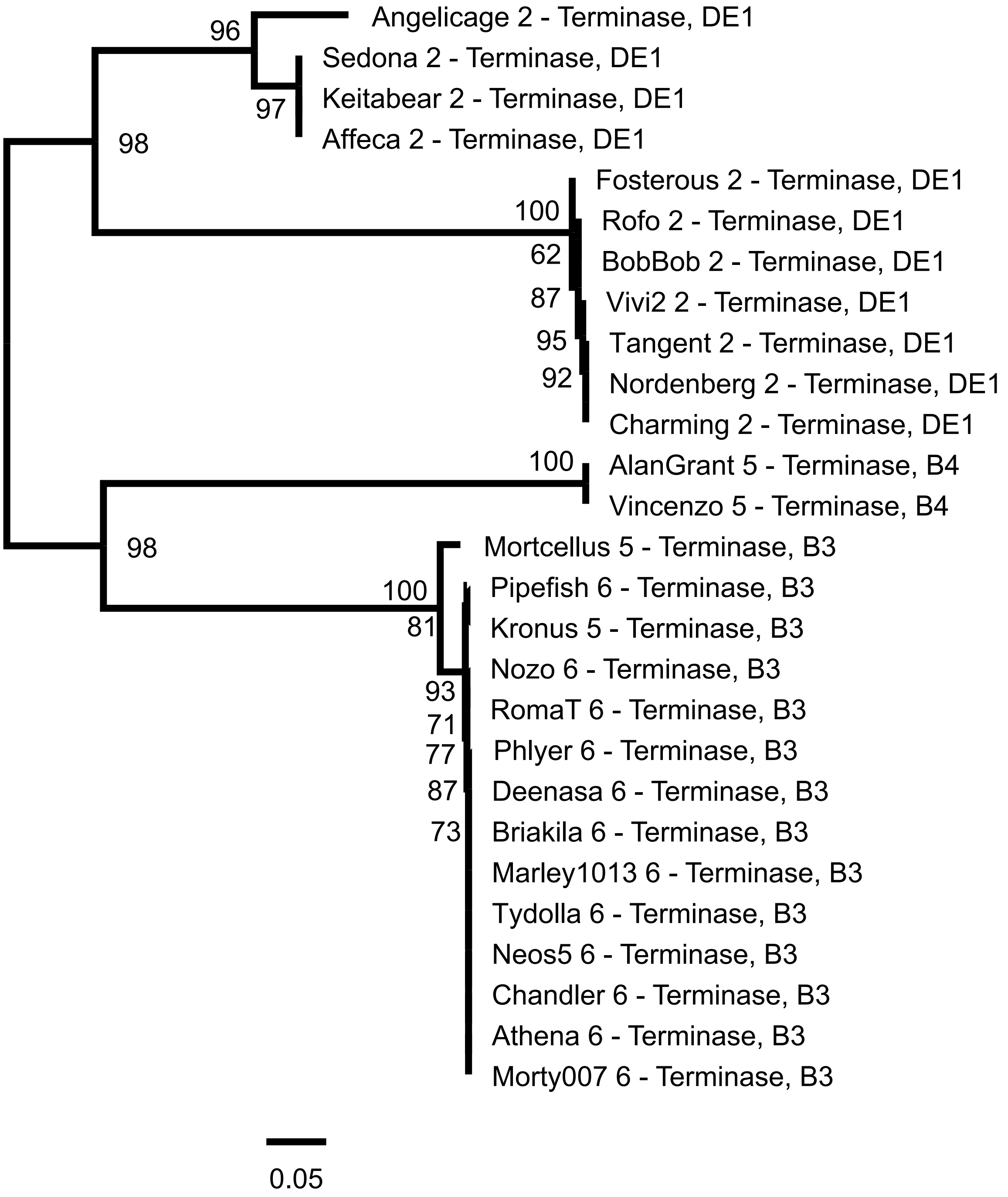


**Supplemental Figure 4**: Maximum likelihood phylogeny of nucleotide sequences encoding inteins with endonuclease-VII-like HEN's using HKY+F+I as the substitution model. Ultrafast bootstrap support is listed on internal nodes. Tip labels include the phage name, gene number, associated extein annotation, and phage cluster. Scale bar measures substitutions per site. Nucleotide sequence alignment is available in the Supplemental Data section under Data for Supplemental Figure 4 – Endonuclease VII Inteins Aligned.fna

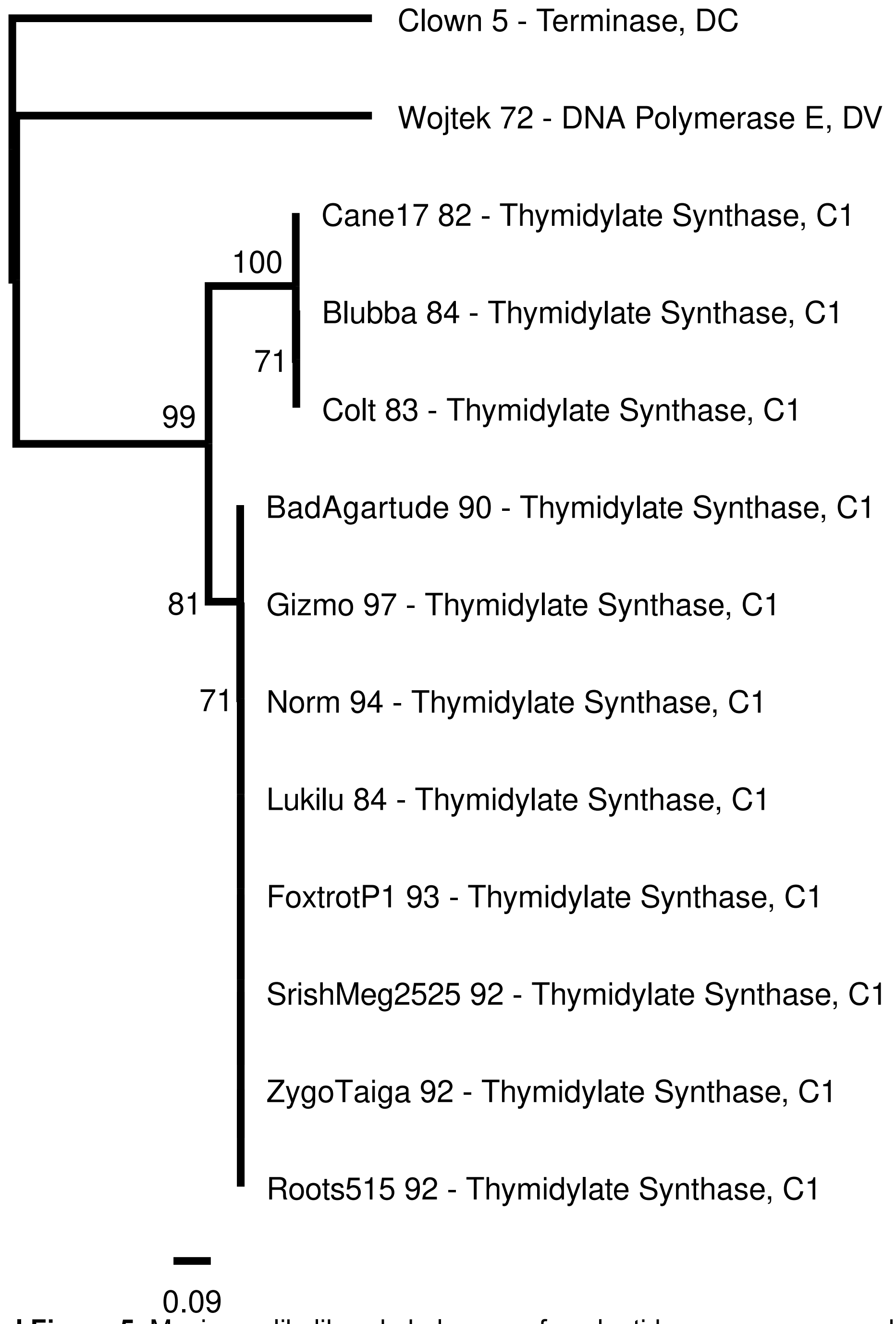


**Supplemental Figure 5**: Maximum likelihood phylogeny of nucleotide sequences encoding inteins with HNH HEN's using TN+F+G4 as the substitution model. Ultrafast bootstrap support is listed on internal nodes. Tip labels include the phage name, gene number, associated extein annotation, and phage cluster. Scale bar measures substitutions per site. Nucleotide sequence alignment is available in the Supplemental Data section under Data for Supplemental Figure 5 – HNH Inteins Aligned.fna

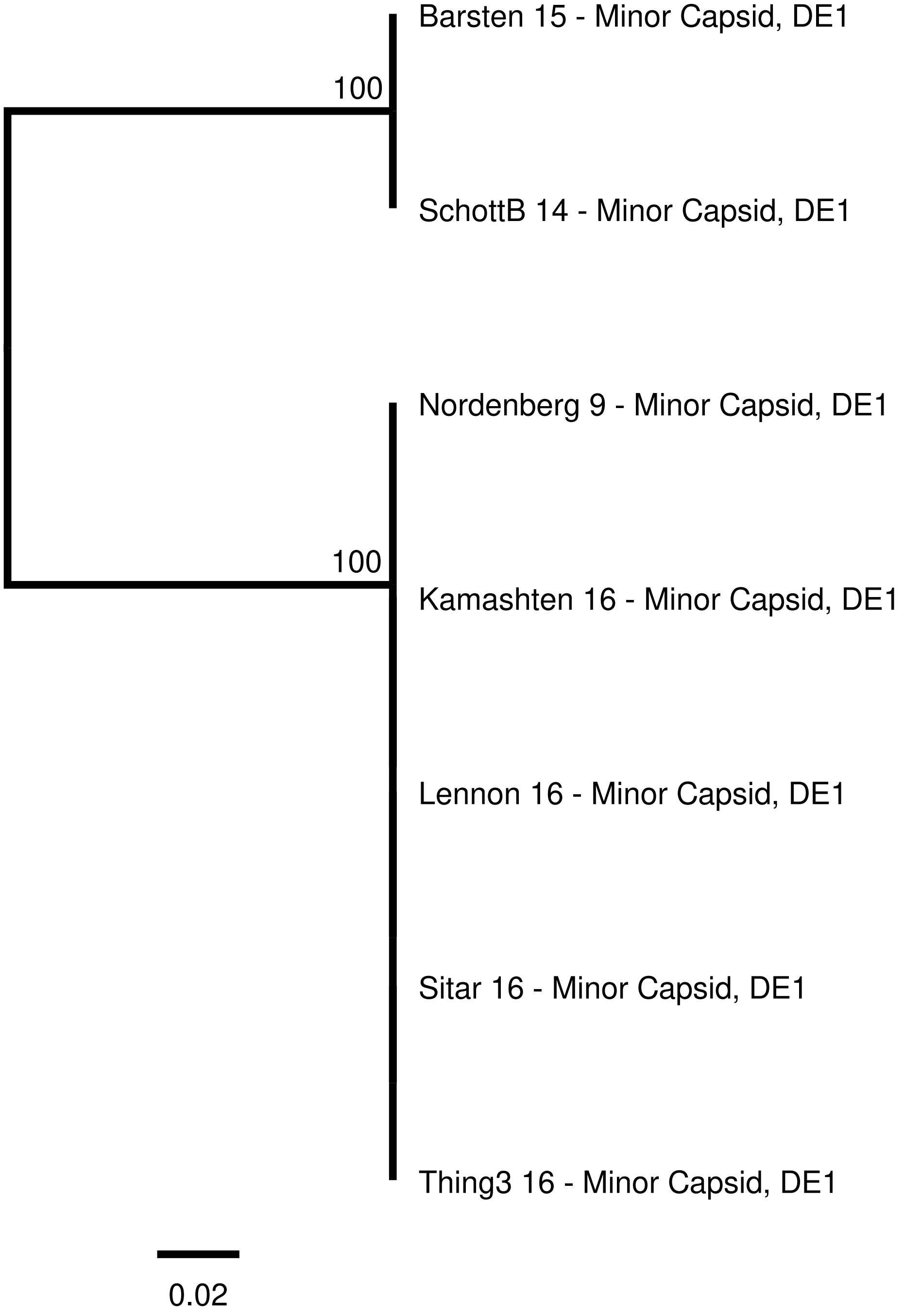


**Supplemental Figure 6**: Maximum likelihood phylogeny of nucleotide sequences encoding inteins with anti-sense LAGLIDADG HEN's using HKY+F as the substitution model. Ultrafast bootstrap support is listed on internal nodes. Tip labels include the phage name, gene number, associated extein annotation, and phage cluster. Scale bar measures substitutions per site. Data for Supplemental Figure 6 – Anti-Sense LAGLIDADG Inteins Aligned.fna

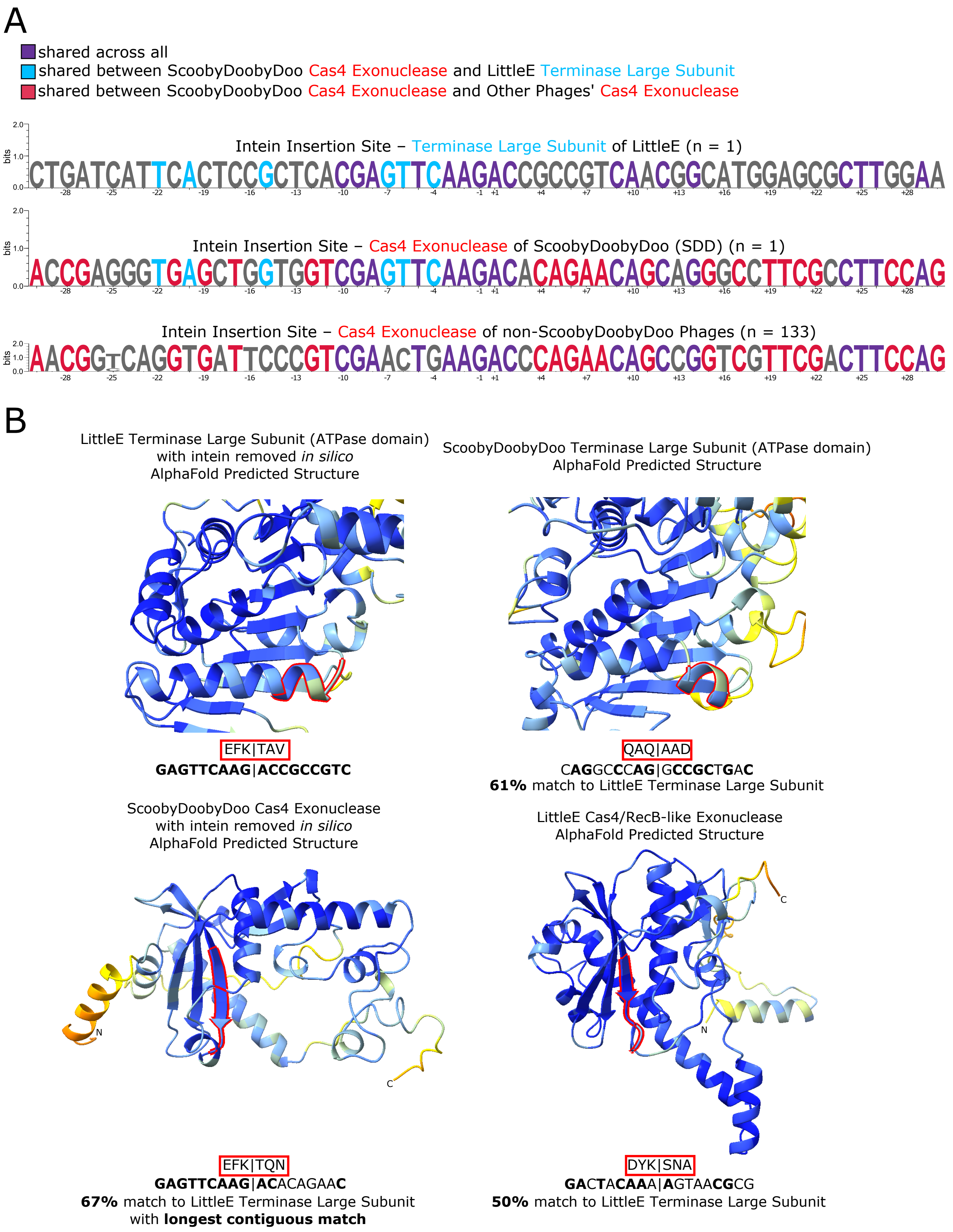
A
shared across all
shared between ScoobyDoobyDoo Cas4 Exonuclease and LittleE Terminase Large Subunit
shared between ScoobyDoobyDoo Cas4 Exonuclease and Other Phages' Cas4 Exonuclease
Intein Insertion Site – Terminase Large Subunit of LittleE (n = 1)
CTGATCATTCACTCCGCTCACGAGTTCAAGACCGCCGTCAACGGCATGGAGCGCTTGGAA
Intein Insertion Site – Cas4 Exonuclease of ScoobyDoobyDoo (SDD) (n = 1)
ACCGAGGGTGAGCTGGTGGTCGAGTTCAAGACACAGAACAGCAGGGCCTTCGCCTTCCAG
Intein Insertion Site – Cas4 Exonuclease of non-ScoobyDoobyDoo Phages (n = 133)
AACGGTCAGGTGATTCCCGTCGAACTGAAGACCCAGAACAGCCGGTCGTTCGACTTCCAG
bits
B
LittleE Terminase Large Subunit (ATPase domain) with intein removed in silico AlphaFold Predicted Structure
ScoobyDoobyDoo Terminase Large Subunit (ATPase domain) AlphaFold Predicted Structure
EFK|TAV
GAGTTCAAG|ACCGCCGTC
QAQ|AAD
CAGGCCCAG|GCCGCTGAC
61% match to LittleE Terminase Large Subunit
ScoobyDoobyDoo Cas4 Exonuclease with intein removed in silico AlphaFold Predicted Structure
LittleE Cas4/RecB-like Exonuclease AlphaFold Predicted Structure
N
C
EFK|TQN
GAGTTCAAG|ACACAGAAC
67% match to LittleE Terminase Large Subunit with longest contiguous match
DYK|SNA
GACTACAAA|AGTAACGCG
50% match to LittleE Terminase Large Subunit

**Supplemental Figure 7**. **A.** Sequence plots were generated using WebLogo3 for the +/-30bp flanking the intein insertion site in the LittleE's terminase large subunit, ScoobyDoobyDoo's Cas4, and the consensus sequence of the other 133 Cas4 sequences with inteins grouping with ScoobyDoobyDoo's. The nucleotides at each position are colored according to the degree to which they are shared across the sequences– a base is colored purple if it is the same across all three sequences, blue if it is shared between ScoobyDoobyDoo Cas4 exonuclease and LittleE terminase large subunit only, or red if it is shared between ScoobyDoobyDoo Cas4 exonuclease and the other Cas4 exonucleases only. **B.** Predicted protein structures were generated for the LittleE terminase large subunit with intein removed (top left), ScoobyDoobyDoo terminase large subunit (top right), ScoobyDoobyDoo Cas4 exonuclease with intein removed (bottom left), and LittleE Cas4/Rec-B like exonuclease (bottom right). The structural regions encoded by the LittleE terminase large subunit intein insertion site and ScoobyDoobyDoo Cas4 intein insertion site are highlighted in red. The +/-9bp encoding these regions is provided, with sequence similarity to the LittleE terminase large subunit intein insertion site indicated. ScoobyDoobyDoo's Cas 4 exonuclease intein insertion site shows the highest degree of sequence similarity to the LittleE terminase large subunit intein insertion site (67%), and longest contiguous stretch of identical nucleotides to the LittleE terminase large subunit intein insertion site. The ScoobyDoobyDoo terminase large subunit region analogous to the LittleE terminase large subunit intein insertion site is 61% identical, and the LittleE Cas4/Rec-B like exonuclease region corresponding to the ScoobyDoobyDoo Cas4 exonuclease intein insertion site is 50% identical.

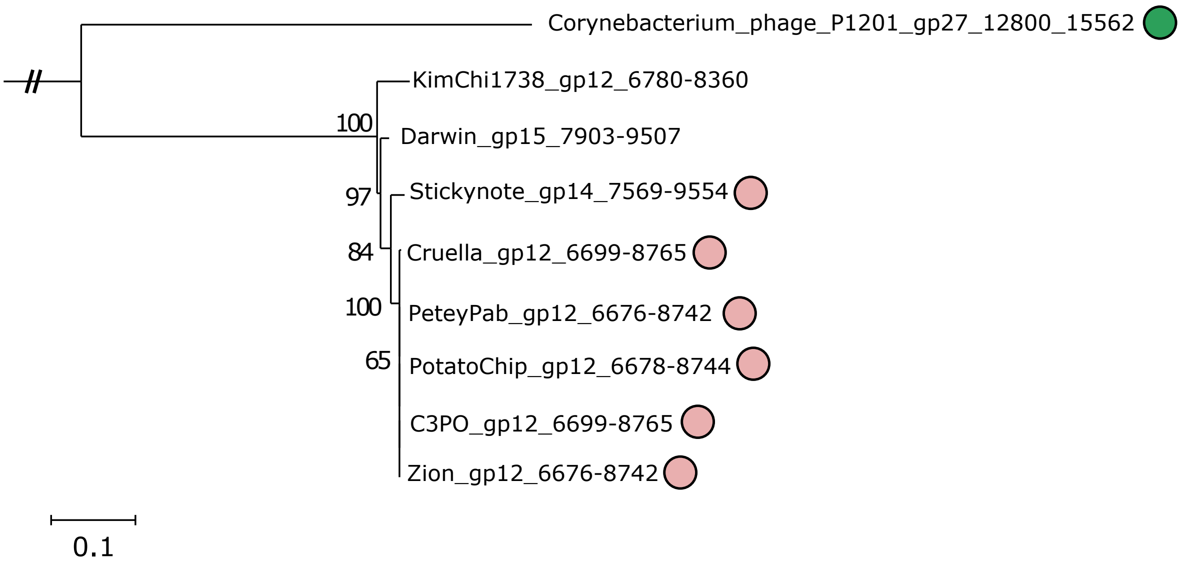


**Supplemental Figure 8**: Subtree (made using GTR+F+R7 as the substitution model) of terminase large subunit coding sequences related to the JonJames terminase large subunit gene. Intein presence is indicated by a circle, with green coloring denoting a complete intein, and the pink denoting a mini intein. The branch connecting to the rest of the terminase phylogeny has been shortened for readability and is indicated by the double slash. Scale bar measures substitutions per site.

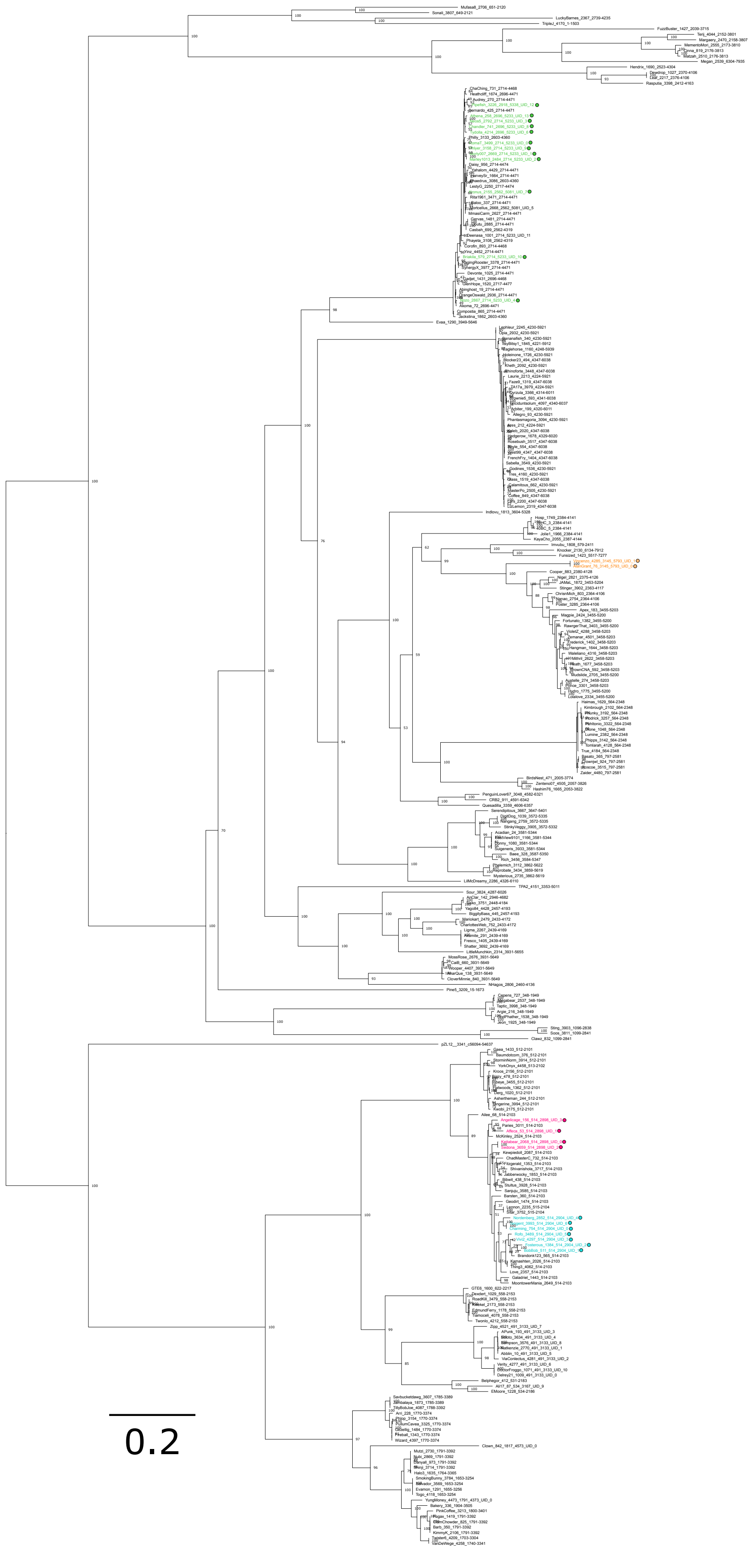
0.2

**Supplemental Figure 9**: Maximum likelihood phylogeny of nucleotide sequences encoding terminase extein sequences with and without an endonuclease-VII-like HEN containing intein. GTR+F+I+R5 was used as the substitution model. Ultrafast bootstrap support is listed on internal nodes. Colored circles indicate the presence of an intein, with the color corresponding to one of the four clans in the intein phylogeny in Supplemental Figure 4. Scale bar measures substitutions per site. The tree image is drawn as a vector graphic that allows one to zoom in to view details.

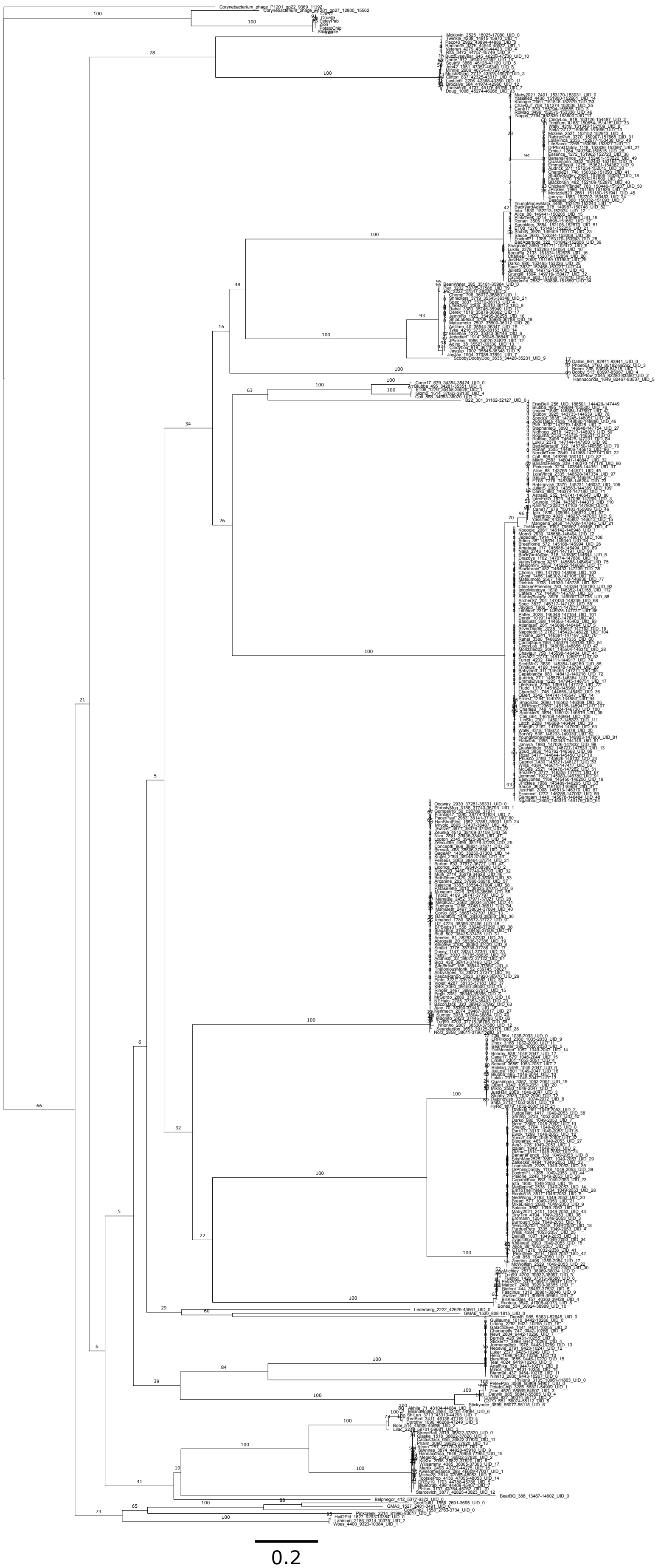

**Supplemental Figure 10**: Un-collapsed version of class 1 intein phylogeny described in Figure 4. Scale bar is 0.1 nucleotide substitutions per site. The tree image is drawn as a vector graphic that allows one to zoom in to view details.

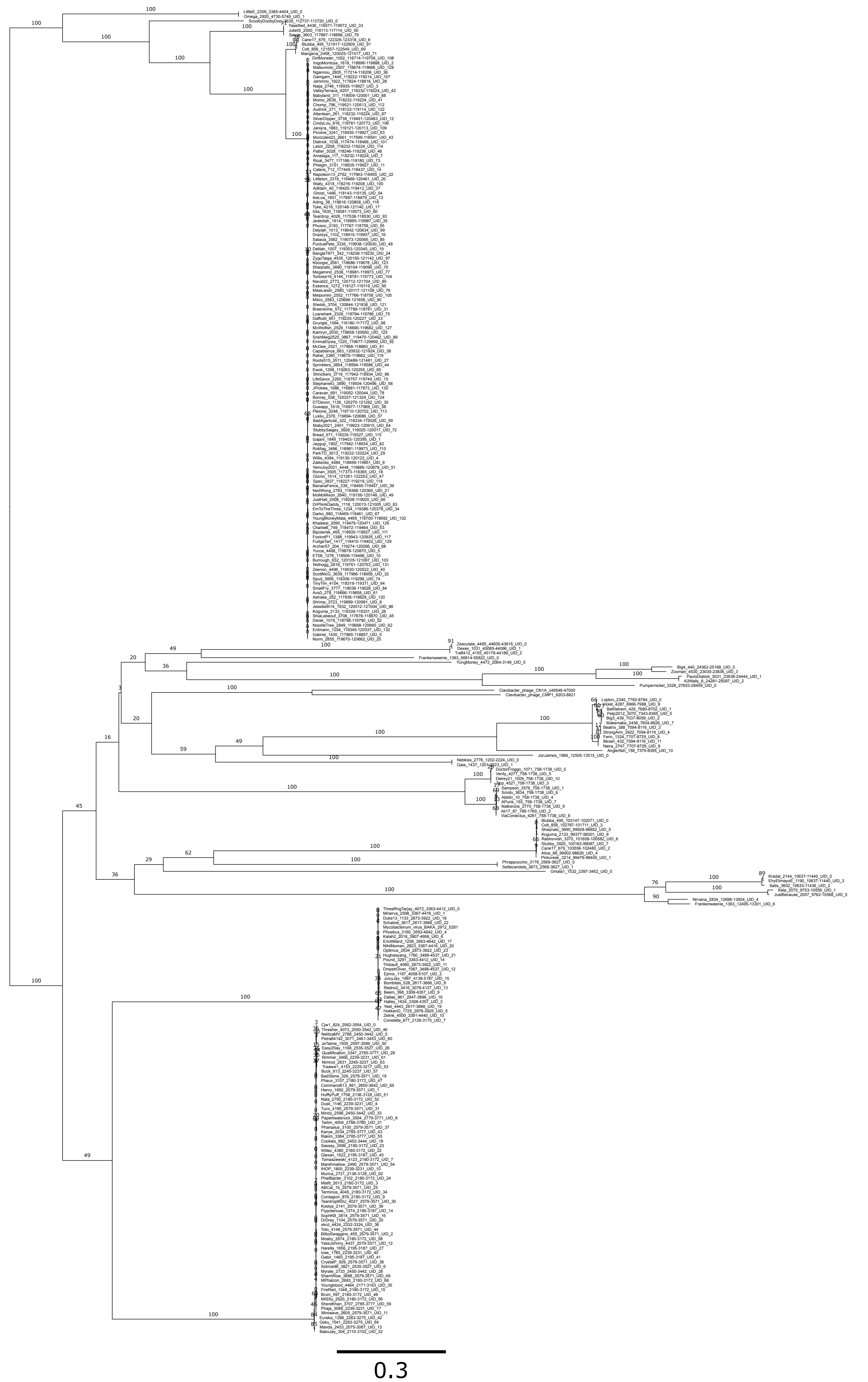
0.3

**Supplemental Figure 11**: Un-collapsed version of class 3 intein phylogeny described in Figure 4. Scale bar is 0.1 nucleotide substitutions per site. The tree image is drawn as a vector graphic that allows to zoom into details.

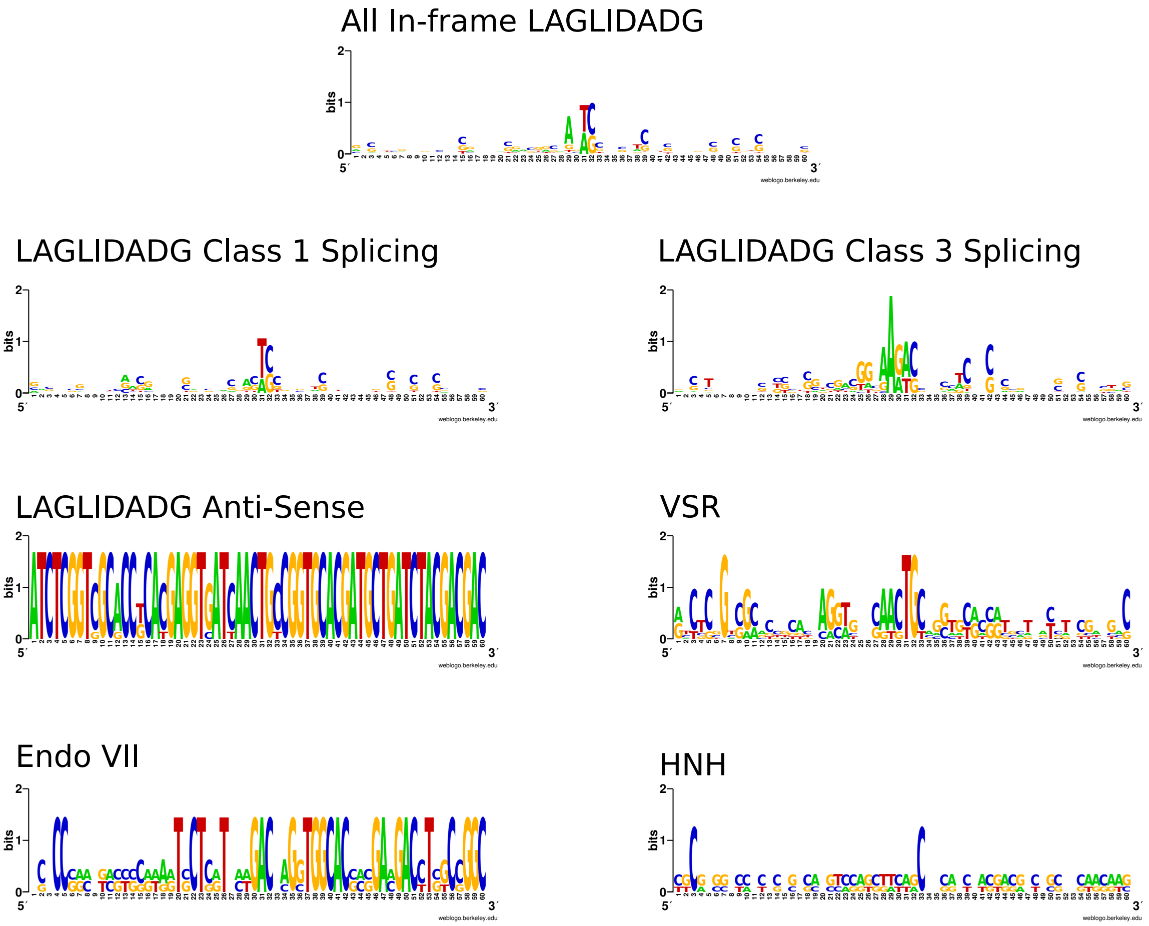


**Supplemental Figure 12**. WebLogo-constructed consensus sequence plots of the intein insertion sites. Each consensus sequence is a subset of the dataset, divided primarily by intein HEN, and further subdivided based on the splicing mechanism for the canonical LAGLIDADG HEN's. Consensus sequences encompass +/-30nt from the site of intein insertion.

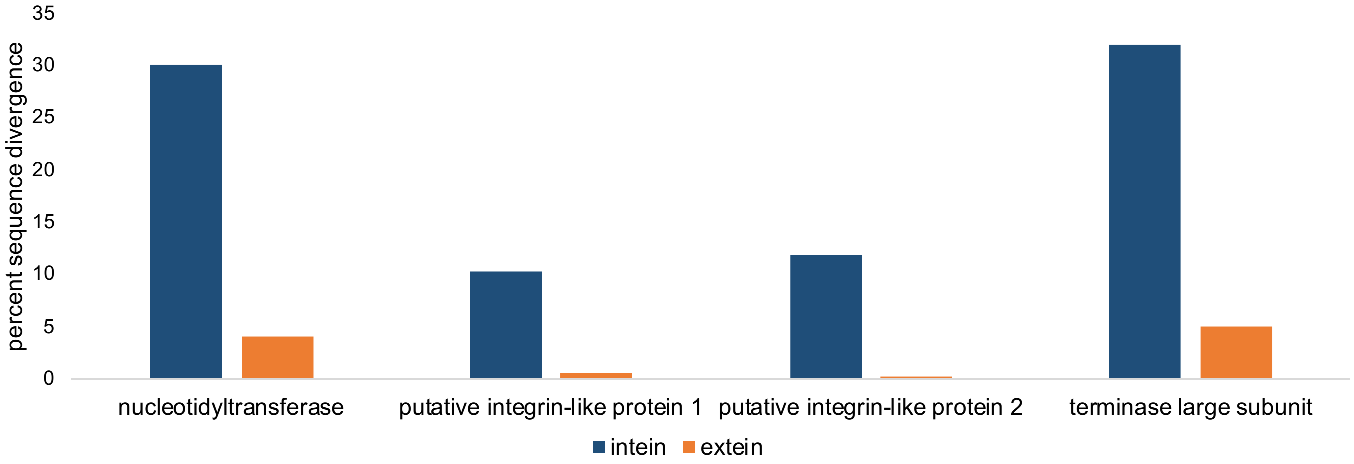


**Supplemental Figure 13.** For Supplemental Results and Discussion section “Sequence Conservation in Inteins and Exteins.” Percent sequence divergence of extein intein pairs. Sequences were aligned using the using the Needleman-Wunsch alignment through the EMBL-EBI Job Dispatcher (Madeira et al. 2024). The following sequences were compared: for nucleotidyltransferase: Bipolarisk_3 and Sebata_3; for putative integrin-like protein 1: CindyLou_262, Ronan_258; putative integrin-like protein 2: CindyLou_262, Melpomini_256; and for the terminase large subunit Violet_10, Anglerfish_12. Intein free homologs from Kamryn_3, Naval_22, and Snazzy_9 were used to define the intein-extein boundaries.

### Supplemental Tables

**Supplemental Table 1**: Table containing all information on all the putative intein sequences within this dataset; including: phage host, CDS #, cluster # (for this study), phage cluster, reviewed gene annotation, class of intein, HEN type, and GenBank Accession if available (all not available on GenBank are currently available on PhagesDB).

[too large to include table directly in document, available as Supplemental_Table1.xlsx]

**Supplemental Table 2.** Results of AU Test run on extein switching events highlighted in class 3 intein phylogeny of Figure 4 (constraining by extein type versus using observed topology) using IQ-TREE

"Event1" = extein switch involving phage Frankenweenie
"Event 2" = extein switch involving phages ScoobyDoobyDoo and LittleE

| **Intein tree topology constraint** | **AU** | **RSS** | **d** | **c** |
|---|---|---|---|---|
| Constrained by Event (Event 1 terminase + Event 1 Cas4) , (Event 2 terminase + Event 2 Cas4) | 0.998 | 3.82 | -2.879 | -0.004 |
| Constrained by gene (Event 1 terminase + Event 2 terminase) , (Event 1 Cas4 + Event 2 Cas4) | 0.002 | 3.82 | 2.879 | 0.004 |

**Supplemental Table 3**: Table containing information on the intein sequence queries used as initial query sequences this study, including: extein descriptions, hosts, start and stop sites, database locations, and accessions when available (or links when not).

| Intein ID | Extein Description | Host | Intein Nucl Start | Intein Nucl End | Database | Accession or Link |
|---|---|---|---|---|---|---|
| AP-Aaphi23 MupF | Putative minor head protein | Haemophilus phage Aaphi23 | N/A | N/A | Inbase | https://inbase.ligsciss.com/iwai/InBase/tools.neb.com/inbase/in-tein841b.html?name=AP-Aaphi23+MupF |
| Hwa MCM-2 | ATP-dependent DNA helicase, CDC | Haloquadratum walsbyi DSM 16790 | N/A | N/A | Inbase | https://inbase.ligsciss.com/iwai/InBase/tools.neb.com/inbase/in-tein7397.html?name=Hwa+MCM-2 |
| Hwa MCM-4 | ATP-dependent DNA helicase, CDC | Haloquadratum walsbyi DSM 16790 | N/A | N/A | Inbase | https://inbase.ligsciss.com/iwai/InBase/tools.neb.com/inbase/in-tein90fe.html?name=Hwa+MCM-4 |
| Pab Lon | ATP-dependent protease LA | Pyrococcus abyssi | N/A | N/A | Inbase | https://inbase.ligsciss.com/iwai/InBase/tools.neb.com/inbase/in-tein16db.html?name=Pab+Lon |
| ArcherS7 CDS 214 | Cas4 Exonuclease | Mycobacterium phage ArcherS7 | 448 | 1440 | Genbank | AGM13864.1 |
| Hma CDC21 | Cell division control protein 21 | Haloarcula marismortui ATCC 43049 | N/A | N/A | Inbase | https://inbase.ligsciss.com/iwai/InBase/tools.neb.com/inbase/in-tein9db8.html?name=Hma+CDC21 |
| Ceu ClpP | ClpP protease | Chlamydomonas eugametos (chloroplast) | N/A | N/A | Inbase | https://inbase.ligsciss.com/iwai/InBase/tools.neb.com/inbase/in-tein9cc2.html?name=Ceu+ClpP |
| Ama MADE823 | Cth-ATCC27405 TerA | Alteromonas macleodii phage | N/A | N/A | Inbase | https://inbase.ligsciss.com/iwai/InBase/tools.neb.com/inbase/in-tein5b4b.html?name=Ama+MADE823 |
| Maeo-N3 Helicase | DEAD/DEAH box helicase domain protein | Methanococcus aeolicus Nankai-3 | N/A | N/A | Inbase | https://inbase.ligsciss.com/iwai/InBase/tools.neb.com/inbase/in-tein53f0.html?name=Maeo-N3+Helicase |
| Mau-ATCC27029 GyrA | DNA gyrase subunit A | Micromonospora aurantiaca ATCC 27029 | N/A | N/A | Inbase | https://inbase.ligsciss.com/iwai/InBase/tools.neb.com/inbase/in-tein64af.html?name=Mau-ATCC27029+GyrA |
| Lsp-PCC8106 GyrB | DNA gyrase subunit B | Lyngbya sp. PCC 8106 | N/A | N/A | Inbase | https://inbase.ligsciss.com/iwai/InBase/tools.neb.com/inbase/inte-inc221.html?name=Lsp-PCC8106+GyrB |
| Cra-CS505 GyrB | DNA gyrase, B subunit | Cylindrospermopsis raciborskii CS-505 | N/A | N/A | Inbase | https://inbase.ligsciss.com/iwai/InBase/tools.neb.com/inbase/in-tein130c.html?name=Cra-CS505+GyrB |
| Alice CDS 175 | DNA Helicase | Mycobacterium phage Alice | 448 | 1524 | Genbank | AEJ94480.1 |
| Bobi CDS 73 | DNA Methylase | Mycobacterium phage Bobi | 592 | 1662 | Genbank | AGS82265.1 |
| Brocaly CDS 61 | DNA Methylase | Mycobacterium phage Brocalys | 2320 | 3312 | Genbank | AMS01768.1 |
| AP-APSE1 dpol | DNA polymerase A (Pol I or family A) | Acyrthosiphon pisum phage | N/A | N/A | Inbase | https://inbase.ligsciss.com/iwai/InBase/tools.neb.com/inbase/in-tein9fce.html?name=AP-APSE1+dpol |
| SaP-SETP12 dpol | DNA polymerase A (Pol I or family A) | Salmonella phage SETP12 | N/A | N/A | Inbase | https://inbase.ligsciss.com/iwai/InBase/tools.neb.com/inbase/in-tein2040.html?name=SaP-SETP12+dpol |
| APMV Pol | DNA polymerase B (alpha or family B) | Acanthomoeba polyphaga Mimivirus | N/A | N/A | Inbase | https://inbase.ligsciss.com/iwai/InBase/tools.neb.com/inbase/in-tein8a5d.html?name=APMV+Pol |
| Hwa PolB-1 | DNA polymerase B (alpha or type-B family) | Haloquadratum walsbyi DSM 16790 | N/A | N/A | Inbase | https://inbase.ligsciss.com/iwai/InBase/tools.neb.com/inbase/in-teina4ec.html?name=Hwa+PolB-1 |
| Hma PolB | DNA polymerase B elongation subunit | Haloarcula marismortui ATCC 43049 | N/A | N/A | Inbase | https://inbase.ligsciss.com/iwai/InBase/tools.neb.com/inbase/in-teina112.html?name=Hma+PolB |
| Cme-boo Pol-II | DNA polymerase II, DP2 subunit, PolC_DP2 | Candidatus Methanoregula boonei 6A8 | N/A | N/A | Inbase | https://inbase.ligsciss.com/iwai/InBase/tools.neb.com/inbase/in-tein517d.html?name=Cme-boo+Pol-II |
| Amax-CS328 DnaX | DNA polymerase III, subunits gamma and tau | Arthrospira maxima CS-328 | N/A | N/A | Inbase | https://inbase.ligsciss.com/iwai/InBase/tools.neb.com/inbase/inte-inc2bd.html?name=Amax-CS328+DnaX |
| Mesp-FS406 PolB-1 | DNA polymerase Pol2,DNA polymerase family B | Methanocaldococcus sp. FS406-22 | N/A | N/A | Inbase | https://inbase.ligsciss.com/iwai/InBase/tools.neb.com/inbase/in-tein4939.html?name=Mesp-FS406+PolB-1 |
| BillKnuckles CDS 54 | DNA Primase | Mycobacterium phage BillKnuckles | 337 | 1272 | Genbank | AER48345.1 |
| EP-Min27 Primase | DNA primase, Helicase | Enterobacteria phage Min27 | N/A | N/A | Inbase | https://inbase.ligsciss.com/iwai/InBase/tools.neb.com/inbase/in-tein6a3e.html?name=EP-Min27+Primase |
| Cmo RPB2 | DNA-directed RNA polymerase beta subunit, fragment 2 | Chlamydomonas moewusii UTEX 97 | N/A | N/A | Inbase | https://inbase.ligsciss.com/iwai/InBase/tools.neb.com/inbase/in-teinae1e.html?name=Cmo+RPB2+%28RpoBb%29 |
| Bde-JEL197 RPB2 | DNA-directed RNA polymerase II, second largest subunit | Batrachochytrium dendrobatidis JEL197 | N/A | N/A | Inbase | https://inbase.ligsciss.com/iwai/InBase/tools.neb.com/inbase/in-teind583.html?name=Bde-JEL197+RPB2 |
| Hwa rPol A | DNA-directed RNA polymerase subunit A | Haloquadratum walsbyi DSM 16790 | N/A | N/A | Inbase | https://inbase.ligsciss.com/iwai/InBase/tools.neb.com/inbase/inte-inc124.html?name=Hwa+rPol+A%27%27 |
| Aeh DnaB-1 | DnaB Helicase | Alkalilimnicola ehrlichei MLHE-1 | N/A | N/A | Inbase | https://inbase.ligsciss.com/iwai/InBase/tools.neb.com/inbase/in-tein3ed9.html?name=Aeh+DnaB-1 |
| Aeh DnaB-2 | DnaB helicase | Alkalilimnicola ehrlichei MLHE-1 | N/A | N/A | Inbase | https://inbase.ligsciss.com/iwai/InBase/tools.neb.com/inbase/in-tein4f00.html?name=Aeh+DnaB-2 |
| Csp-PCC7425 DnaB | DnaB helicase | Cyanothece sp. PCC 7425 | N/A | N/A | Inbase | https://inbase.ligsciss.com/iwai/InBase/tools.neb.com/inbase/in-teinf9ca.html?name=Csp-PCC7425+DnaB |
| Gob DnaE | DnaE pol subunit, DNA polymerase III alpha subunit | Gemmata obscuriglobus UQM2246 | N/A | N/A | Inbase | https://inbase.ligsciss.com/iwai/InBase/tools.neb.com/inbase/in-tein3f38.html?name=Gob+DnaE |
| Mcht-PCC7420 DnaE-1 | DnaE pol subunit, DNA polymerase III alpha subunit | Microcoleus chthonoplastes PCC7420 | N/A | N/A | Inbase | https://inbase.ligsciss.com/iwai/InBase/tools.neb.com/inbase/in-teinf5cb.html?name=Mcht-PCC7420+DnaE-1 |
| Pma-ExH1 DnaE | DnaE pol subunit, DNA polymerase III alpha subunit | Persephonella marina EX-H1 | N/A | N/A | Inbase | https://inbase.ligsciss.com/iwai/InBase/tools.neb.com/inbase/in-tein3e0e.html?name=Pma-ExH1+DnaE |
| Bde-JEL423 eIF-5B | Eukaryotic translation initiation factor 5B | Batrachochytrium dendrobatidis JEL423 | N/A | N/A | Inbase | https://inbase.ligsciss.com/iwai/InBase/tools.neb.com/inbase/in-tein0ac5.html?name=Bde-JEL423+eIF-5B |
| Fac-Fer1 SufB | FeS assembly protein SufB | Ferroplasma acidarmanus | N/A | N/A | Inbase | https://inbase.ligsciss.com/iwai/InBase/tools.neb.com/inbase/in-tein01d8.html?name=Fac-Fer1+SufB+%28Fac+Pps1%29 |
| Pfl Fha BIL | Filamentous hemagglutinin | Pseudomonas fluorescens Pf-5 | N/A | N/A | Inbase | https://inbase.ligsciss.com/iwai/InBase/tools.neb.com/inbase/in-tein42c5.html?name=Pfl+Fha+BIL |
| Dhan GLT1 | Glutamate synthase | Debaryomyces hansenii CBS767 | N/A | N/A | Inbase | https://inbase.ligsciss.com/iwai/InBase/tools.neb.com/inbase/in-teina2a5.html?name=Dhan+GLT1 |
| MP-Be gp51 | gp51 | Mycobacteriophage Bethlehem | N/A | N/A | Inbase | https://inbase.ligsciss.com/iwai/InBase/tools.neb.com/inbase/in-tein8a26.html?name=MP-Be+gp51 |
| Cce Hyp1-Csp-2 | Hypothetical, UPF0027-containing protein | Cyanothece sp. ATCC 51142 | N/A | N/A | Inbase | https://inbase.ligsciss.com/iwai/InBase/tools.neb.com/inbase/in-teina551.html?name=Cce+Hyp1-Csp-2 |
| Mja KlbA | KlbA, kilB operon ORF A | Methanococcus jannaschii | N/A | N/A | Inbase | https://inbase.ligsciss.com/iwai/InBase/tools.neb.com/inbase/in-tein5077.html?name=Mja+KlbA |
| Mesp-FS406-22 LHR | Large helicase related protein | Methanocaldococcus sp. FS406-22 | N/A | N/A | Inbase | https://inbase.ligsciss.com/iwai/InBase/tools.neb.com/inbase/in-teina3f0.html?name=Mesp-FS406-22+LHR |
| Alice CDS 3 | Nucleotidyltransferase | Mycobacterium phage Alice | 577 | 1581 | Genbank | AEJ94269.1 |
| Cwa PEP | Phosphoenolpyruvate synthase | Crocosphaera watsonii WH 8501 | N/A | N/A | Inbase | https://inbase.ligsciss.com/iwai/InBase/tools.neb.com/inbase/in-tein90b6.html?name=Cwa+PEP |
| Aeneas CDS 53 | Phosphoesterase | Mycobacterium phage Aeneas | 283 | 1233 | Genbank | AFL48067.1 |
| Beanwater CDS 82 | Portal Protein | Mycobacterium phage BeanWater | 346 | 1149 | Genbank | ATN87536.1 |
| Maeo-N3 RtcB | Protein of unknown function UPF0027 | Methanococcus aeolicus Nankai-3 | N/A | N/A | Inbase | https://inbase.ligsciss.com/iwai/InBase/tools.neb.com/inbase/in-tein2786.html?name=Maeo-N3+RtcB |
| Abr PRP8 | PRP8, pre-mRNA splicing factor | Aspergillus brevipes FRR2439 | N/A | N/A | Inbase | https://inbase.ligsciss.com/iwai/InBase/tools.neb.com/inbase/in-tein37bf.html?name=Abr+PRP8 |
| Alice CDS 247 | Putative Integrin | Mycobacterium phage Alice | 562 | 1326 | Genbank | AEJ94469.1 |
| Pho RadA | RadA DNA repair protein | Pyrococcus horikoshii OT3 | N/A | N/A | Inbase | https://inbase.ligsciss.com/iwai/InBase/tools.neb.com/inbase/in-tein1634.html?name=Pho+RadA |
| Mbo RecA | RecA | Mycobacterium bovis subsp. bovis AF2122/97 | N/A | N/A | Inbase | https://inbase.ligsciss.com/iwai/InBase/tools.neb.com/inbase/inte-inc883.html?name=Mbo+RecA |
| Mch RecA | RecA | Mycobacterium chitae | N/A | N/A | Inbase | https://inbase.ligsciss.com/iwai/InBase/tools.neb.com/inbase/in-tein86ca.html?name=Mch+RecA |
| Hwa RCF | Replication factor C | Haloquadratum walsbyi DSM 16790 | N/A | N/A | Inbase | https://inbase.ligsciss.com/iwai/InBase/tools.neb.com/inbase/in-tein58a6.html?name=Hwa+RCF |
| Maeo RFC | Replication factor C | Methanococcus aeolicus Nankai-3 | N/A | N/A | Inbase | https://inbase.ligsciss.com/iwai/InBase/tools.neb.com/inbase/inte-inc963.html?name=Maeo+RFC |
| Mein-ME RFC | Replication factor C | Methanocaldococcus infernus ME | N/A | N/A | Inbase | https://inbase.ligsciss.com/iwai/InBase/tools.neb.com/inbase/in-tein2680.html?name=Mein-ME+RFC |
| Mja r-Gyr | Reverse gyrase | Methanococcus jannaschii | N/A | N/A | Inbase | https://inbase.ligsciss.com/iwai/InBase/tools.neb.com/inbase/in-tein5839.html?name=Mja+r-Gyr |
| Ace RIR1 | Ribonucleoside diphosphate reductase, class I | Acidothermus cellulolyticus 11B | N/A | N/A | Inbase | https://inbase.ligsciss.com/iwai/InBase/tools.neb.com/inbase/in-tein6bce.html?name=Ace+RIR1 |
| Avin RIR1 BIL | Ribonucleoside- diphosphate reductase, alpha subunit | Azotobacter vinelandii | N/A | N/A | Inbase | https://inbase.ligsciss.com/iwai/InBase/tools.neb.com/inbase/in-teinfc10.html?name=Avin+RIR1+BIL |
| BsuP-M1918 RIR1 | Ribonucleoside- diphosphate reductase, alpha subunit | B.subtilis M1918 phage | N/A | N/A | Inbase | https://inbase.ligsciss.com/iwai/InBase/tools.neb.com/inbase/in-teinfdd0.html?name=BsuP-M1918+RIR1 |
| Gvi RIR1-1 | Ribonucleoside- diphosphate reductase, alpha subunit | Gloeobacter violaceus, PCC 7421 | N/A | N/A | Inbase | https://inbase.ligsciss.com/iwai/InBase/tools.neb.com/inbase/in-tein6d14.html?name=Gvi+RIR1-1 |
| Hwa RIR1-2 | Ribonucleoside- diphosphate reductase, alpha subunit | Haloquadratum walsbyi DSM 16790 | N/A | N/A | Inbase | https://inbase.ligsciss.com/iwai/InBase/tools.neb.com/inbase/in-teina5c6.html?name=Hwa+RIR1-2 |
| Pab RIR1-1 | Ribonucleoside- diphosphate reductase, alpha subunit | Pyrococcus abyssi | N/A | N/A | Inbase | https://inbase.ligsciss.com/iwai/InBase/tools.neb.com/inbase/in-tein7ae9.html?name=Pab+RIR1-1 |
| Aave-AAC001 RIR1 | ribonucleoside-diphosphate reductase, class I | Acidovorax avenae subsp. citrulli AAC00-1 | N/A | N/A | Inbase | https://inbase.ligsciss.com/iwai/InBase/tools.neb.com/inbase/in-tein70bc.html?name=Aave-AAC001+RIR1 |
| Mcht-PCC7420 RIR1-1 | ribonucleoside-triphosphate reductase | Microcoleus chthonoplastes PCC 7420 | N/A | N/A | Inbase | https://inbase.ligsciss.com/iwai/InBase/tools.neb.com/inbase/inte-incb99.html?name=Mcht-PCC7420+RIR1-1 |
| CbP-C-St RNR | Ribonucleotide reductase, class III | Clostridium botulinum phage C-St | N/A | N/A | Inbase | https://inbase.ligsciss.com/iwai/InBase/tools.neb.com/inbase/in-teinad95.html?name=CbP-C-St+RNR |
| Mja RNR-2 | Ribonucleotide reductase, class III | Methanococcus jannaschii | N/A | N/A | Inbase | https://inbase.ligsciss.com/iwai/InBase/tools.neb.com/inbase/in-tein249f.html?name=Mja+RNR-2 |
| Maeo RNR | Ribonucleotide-triphosphate reductase, class III | Methanococcus aeolicus Nankai-3 | N/A | N/A | Inbase | https://inbase.ligsciss.com/iwai/InBase/tools.neb.com/inbase/in-tein8576.html?name=Maeo+RNR |
| Dra Snf2-c | SNF2/Rad54 helicase | Deinococcus radiodurans R1 TIGR | N/A | N/A | Inbase | https://inbase.ligsciss.com/iwai/InBase/tools.neb.com/inbase/inte-ine00e.html?name=Dra+Snf2-c |
| Alice CDS 239 | Terminase | Mycobacterium phage Alice | 865 | 1671 | Genbank | AEJ94461.1 |
| ABCat CDS 8 | Terminase, Large Subunit | Mycobacterium phage ABCat | 277 | 1269 | Genbank | GR48848.1 |
| AlanGrant CDS 5 | Terminase, Large Subunit | Mycobacterium phage AlanGrant | 787 | 1686 | Genbank | AKF14670.1 |
| PauloDiaboli CDS 72 | Terminase, Large Subunit | Microbacterium phage PauloDiaboli | 295 | 1101 | Genbank | QIG57807.1 |
| Cpa ThrRS | Threonyl-tRNA synthetase | Candida parapsilosis CLIB214 | N/A | N/A | Inbase | https://inbase.ligsciss.com/iwai/InBase/tools.neb.com/inbase/in-tein29cd.html?name=Cpa+ThrRS |
| Gizmo CDS 97 | Thymidylate Synthase | Mycobacterium phage Gizmo | 253 | 1023 | Genbank | AGM13387.1 |
| Hut MCM-2 | transcriptional regulator, XRE family | Halorhabdus utahensis DSM 12940 | N/A | N/A | Inbase | https://inbase.ligsciss.com/iwai/InBase/tools.neb.com/inbase/in-tein0802.html?name=Hut+MCM-2 |
| Maeo-N3 UDP GD | UDP-glucose 6-dehydrogenase | Methanococcus aeolicus Nankai-3 | N/A | N/A | Inbase | https://inbase.ligsciss.com/iwai/InBase/tools.neb.com/inbase/in-tein9b7a.html?name=Maeo-N3+UDP+GD |
| Cgl VMA | Vacuolar ATPase subunit A | Candida glabrata | N/A | N/A | Inbase | https://inbase.ligsciss.com/iwai/InBase/tools.neb.com/inbase/in-tein5b26.html?name=Cgl+VMA |
| Pab VMA | Vacuolar ATPase subunit A | Pyrococcus abyssi | N/A | N/A | Inbase | https://inbase.ligsciss.com/iwai/InBase/tools.neb.com/inbase/in-tein0466.html?name=Pab+VMA |

**Supplemental Table 4**: Table summarizing the results from searching the Invader-Sim simulated mixed intein and non-intein containing sequence databases to test ICE-BLAST. Summary plots for this data can also be found in the zip file: Supplemental_Material_Invader_Sim_Results_Graphs.zip.

[too large to include table directly in document, available as Supplemental_Table4.xlsx]

**Supplemental Table 5:** Intein insertion site consensus sequence (at 60%) for each of the clusters created for this study.

| Cluster # | Centroid Phage | 60% Consensus Insertion Sequence (+/-30) |
|---|---|---|
| 0 | P1201 | CTCAAGAACATCCTCCGTCAGAAGCATCACTCTGTGCTGGAGCACACCAGCTTCACGTTC |
| 1 | LRRHood | CCNGAGCTGATCGAGGCCTACGGNTGGGATACCAAGACCGGCTATCACGCGCTGCGCCTG |
| 2 | Lahirium | GAGTACGACTACCAGGTCGAGTACTTGGAGTCGCAAATGCGTGGCGCAGACTTTGAGCGC |
| 3 | Pinkcreek | GAGGTGATGGAGAATTGGAGGTACATCGACTCCTTCGGCCTGGGATCCAACGTCGGACGC |
| 4 | Job42 | CAGAAACGCCGCAACGTTCAAATGCACGTGTGCCCCTTGCAGTTCGACATCGTTGACCGG |
| 5 | Belphegor | CTCGAGGGCGCCAGCGGCATGAATCACTGGTCGGCGTGGTCGCTCGACGAGTCCGAGATC |
| 6 | Darwin | GCCGTCCTAACCAACATTTCCTTCATGGAGTGCGTCCATGCACGCTCCTACTCCTCAATC |
| 7 | Magnito | TGGATCGGGGCTCATGAAGGGAACCACGACTCCCGGGCCAGGGACTATCTCTCCAAGAAC |
| 8 | Settecandela | CCGTCCTCACCCTGGCAGAAGTCCGGCAAGTGCTACGGCCTCTACCAGGAGGGCTCGGTG |
| 9 | YungMoney | ATGATCTTCATCGGCCCCCAAACCGGCAAGAGCACCCGCACCTCACGCTGGTTGCCGTTC |
| 10 | Kamryn | AACGGTCAGGTGATTCCCGTCGAACTGAAGACCCAGAACAGCCGGTCGTTCGACTTCCAG |
| 11 | CN1A | GCCATCAAGAGCGCCAACGGCTGCGGGAAGAGCATGGACCTCGCAGACCTCATCACTTGG |
| 12 | ZygoTaiga | CGCGAGCATCACCGTCACAGGTCGGGCTTTAGCTACAACGAGGAGAGTGGGCGCTACAAG |
| 13 | C3PO | AACGTNGGNAGAATCTCNATGTCCAACTTNTGTTCGGAGATTCTACAGCCTCAGACGGCA |
| 14 | GordDuk1 | GAAACCTCTGTCGAAGGTTATAACGTACTTTGCGGGATCATTGACGAGGGTGACAGCCAT |
| 18 | BobBob | TCACCGACCTACCGCGAAGTGCTCGTNATGACAAGGTGGCACGAGGACGACCTCGCCGGC |
| 19 | ChaylaJr | CCGTGGCACAAGCAGGGTCGCAAGGGGTACTGCATCTTCATCGGNGACGAGAAGCCCTAC |
| 20 | JayJay | GAAGAAGAGCTCAAGACCATCAGACACTGGTGCCGCCTCTTCTATGCCACCCACGACCTG |
| 35 | ShiLan | ATCCGCGACTCNATCGCNTGGCTNTACGGGAGCGGCTTCCCGAAGTCGCTCGACGTNTCN |
| 36 | BlueCrab | ATCCGNGACTCCATCGCNTGGCTGTACGGCAGCGGGTTCCCGAAGTCGCTCGACGTATCC |
| 37 | Petp2012 | AGCCACACGGTAGCTATCGCCGAGGGAGAGTGCGACACGATTGCGGCGTCTGTCGCGGGG |
| 38 | Kalah2 | GGCCTGATGGTTAGCCGGCAGAACGGCAAGAGCATCCTGGAGGCCATTGAGCTCGCTGGG |
| 39 | Cantare | GACAAGGAAAACGACAAGTACAAGCTACAGAGTGCGGAATATCAGTTCATTGGCTTCGAC |
| 40 | JonJames | GGTAAGGCCACCCTTGTGCTAGCAGATGAAACCCACCTCTGGAAGACTAACGAGCTCCAC |
| 41 | Command613 | TGGATTGAGCTCGCGCGCAAGAACGGCAAAACCGAACTCCTCGCCGGAATCATGCTCTAC |
| 42 | ViaConlectus | ATGATCTTCACNCCGTCGCAGGTCGGCAAGTCGACGCGTGTGTCGCAGTGGTTCCCGTTT |
| 43 | Malisha | GCCTCTGCGAACTTTGGTTCGCACGACAACTGTAAGTGTGTGGCTGTTCCCGCTTTTGGC |
| 44 | Sejanus | GACGCCGACGAGCTGCAGAAAGTGTGGGTGTGCACCATCGACGGCAAGACCAGGCCATCG |
| 46 | Cece | GACGTGCGAGAGGTTATCGTTGAAGGTGAATCCGGTATTCTCGCCGTCTCGCCCGAGTCC |
| 47 | Sedona | TCGCCGACNTACCGCGAAGTGCTCGTTATGACGCGCTGGCACGAGGACGACCTCGCCGGC |
| 48 | Astraea | GCCATGACTTACCTGGCATCGAGCCCCTACTCGAANATNGGCAAGTTCTACGAGCTCTAC |
| 53 | BearBQ | GTGTGGGGTGGTGTGAACTGGGGGGACTGCTCCGGAGCCATGTCGGCGTTCGCGCGTATG |
| 54 | Gizmo | AAGCTGCTCATCCCGGAGATCAAGTTCTCCACGCTCAACCTGCGTGACGAGGCCCAAGAG |
| 55 | GordTnk2 | GAGAAAGATTATGCCGAAGAGGCTTGGAATACTATTCACGGTCGTATTGCTAGCCGTTTT |
| 56 | Gmala1 | ATGTTCACTGGCGGCATTGGCGTCGGGAAATCAACCCTAGCCAGTATCGCGTTGACCTAT |
| 57 | ScoobyDoobyDoo | ACCGAGGGTGAGCTGGTGGTCGAGTTCAAGACACAGAACAGCAGGGCCTTCGCCTTCCAG |
| 58 | Anglerfish | ATCACGATCGCCGCGGTCTCCCAGGACCAGACGAAGAACACGTTCTCGCTGTTCCCGATC |
| 59 | CMP1 | ATCGTAAAGTCTTGCAACGGTGTCGGCAAGACGCGCCTGGCCGGAGACCTCGTGACGTGG |
| 60 | Vivi2 | ATCTCGGTCGCACCNCACGAGGTGATCAACTGCCGGTGCACGATGCTNATNTACGACGAC |
| 62 | PauloDiaboli | GACGTNCGAGAGGTTATCGTTGANGGTGAATCGGGCATCNTCGCGGTCTCNCCGNNNTCC |
| 63 | Phlyer | AGCCCCGAGGCGTCCATGATCCTGATCCAGACCAGGTGGCACCCCGAAGACCTGTCCGGC |
| 69 | Colt | CCCGAGCTGATCGCCAAGCATGGCTGGGATACCAAGACCGGCTATCACGCGCTGCGCCTG |
| 70 | B22 | CCGGTGGATGTGATCTGTGGCGGGAGCCCCTGTCAGGACCTGTCCATGGCCGGACGGCGG |
| 71 | GMA3 | GAGACCTCGGTGGAGGGTTATAACGTTCTCTGCGGGATCATTGACGAGGGCGATAGCCAT |
| 72 | Scioto | ATCTCGGTGGCGCCGCACGAGGTGATCAACTGCCGGTGCACGATGCTGATCTACGACGAC |
| 73 | Phoebus | GAAATCGTGATCACCCAGAAGCTCCACGGCTCGTCTATCCGTGTTGGGCGAGTGCCTTGC |
| 74 | Kela | GTCGTCGCCTCCTGCCACGGCACCGGNAAGTCGATGATCGCCTCGGTNCTCGCNTGCTGG |
| 79 | Lederberg | ATCCCCGCAGCCGTTGGCGTCGACATCGGCTGCGGCATGATCGCCGCCCGCACCCGCTAC |
| 80 | Nebkiss | GACCCCGCCAGTTTTATTATCTTGAATGAGAGTCACCACATGACCGCCACTAATGGTGGG |
| 81 | Traft412 | GACGGTGAGGTGCTGGTGGTCGACTATAAAACGGGACTCAAGCCCGGAGATGACTTTCAG |
| 82 | REQ3 | GCCCGCGCCGCAGGGTTTCAGCATCCGAACTGCGGCCATGCCGTCAGCGCGTTCGTGCCC |
| 83 | Sticker17 | GCAGGTGGATTTAACCAGGGGGTTGCTCACACACAGGTATATCTGTGGATGCTCAACGCT |
| 84 | Philon9 | GGTTCCATCTACAAGGTACCGGTGGAGAACTCTTTGTGGTACCCGCTGCTGATGCAGAAC |
| 87 | GMA6 | CAGCGCGACGCAGACAAGTACAAGTTCCAGTCCGCTGAGTATCAGTTCATCGGCTTCGAT |
| 88 | Omega | CTGATCATTCACTCCGCTCACGAGTTCAAGACCGCCGTCAACGGCATGGAGCGCTTGGAA |
| 89 | Shaqnato | TTCCTGAACTGGTCGGCGGGCTCGGGGAAAAGCTTTTGTGCCGCCGCCGGGGCCCGCGCG |
| 90 | Frankenweenie | CCACGCAAGAAGACCCTGGTCGACTGGAAGACAACCGACCTGAAGAAGCTGAAGAAGTAC |
| 91 | Kewpiedoll | ATTTCGGTCGCACCTCACGAGGTGATCAACTGCAGGTGCACCATGCTTGTGTACGACGAC |
| 92 | Pumpernickel | GAAGTGCGAGAGGTTATCGTTGAAGGTGAATCTGGAATCCTCGCTATCTCCCCGCCCGAT |
| 102 | P1201 | AACCTCTCCCTACCCAAGGGTTCCGCAAAGACTGAGTTGATGGCGCTGATCTCGAACGTC |
| 103 | P1201 | AACATTGGTCGCATTAAGAGCTCCAACTTGTGCTCGGAGATTTTCCAGCCATCCTCGCCT |
| 104 | Mcklovin | GACTACGGCGCCGGCACCAACGCCGACCACTCGGACCACGTNCACTGGGCGATGGCNGCG |
| 105 | Vincenzo | GCCCCCGATGCGTCGATAATCCTCATTCAGACGAGGTGGCACCCCGAAGACTTGGCGGGC |
| 106 | Wojtek | TTCATGGTCACTTCGGCGACCTACAGCTTCAACATCGCTCACTGCGTCAGCTACAGCATG |
| 107 | Nordenberg | ATCTCGGTCGCACCNCACGAGGTGATCAACTGCCGGTGCACGATGCTGATCTACGACGAC |
| 108 | Clown | CGCGGCGGCCTCACCGGCCAGCCCATGGACTGCGGGATCATCGACGACCCCGTGAAGGAC |

**Supplemental Table 6**. For Supplemental Results and Discussion section “Sequence Conservation in Inteins and Exteins.” Parameters describing strength of selection estimated from the alignments of the nucleotidyltransferase inteins and exteins.

| | Shape parameter for the Gamma distribution to describe Among Site Rate variation [$] | Fraction of sites estimated to be under purifying selection (dN/dS<1) [$] | Average dN/dS value for sites under purifying selection[$] |
|---|---|---|---|
| Intein | 4.7 [1.92, 8.01] | 0.740 [0.653, 0.833] | 0.0777 [0.004, 0.185] |
| Extein | 0.065 [0.013, 0.095] | 0.895 [0.857, 0.934] | 0.0066 [0.0012, 0.0133] |

[$] The 95% High Probability Density interval is given in square brackets.

**Supplemental Data Descriptions**

**The following are all stored in Supplemental Data.zip:**

- **Identified Inteins - Extein Nucleotide Sequences.zip:** Folder containing the extein only (inteins removed *in silico*) sequences of all 784 inteins, organized by intein cluster. The filenames indicate the type of host protein each cluster of inteins is found in.
- **Identified Inteins - Extein With Intein Nucleotide Sequences.zip:** Folder containing the extein with intein sequences of all 784 inteins, organized by intein cluster. The filenames indicate the type of host protein each cluster of inteins is found in.
- **Identified Inteins - Intein Nucleotide Sequences.zip:** Folder containing the intein only (extein removed *in silico*) sequences of all 784 inteins, organized by intein cluster. The filenames indicate the type of host protein each cluster of inteins is found in.
- **Data for Supplemental Figure 3 – Minor Capsid VSR Inteins Aligned.fna** Nucleotide sequence alignment of identified VSR-containing inteins, used to generate phylogeny in Supplemental Figure 3.
- **Data for Supplemental Figure 4 – Endonuclease VII Inteins Aligned.fna** Nucleotide sequence alignment of identified endonuclease VII-containing inteins, used to generate phylogeny in Supplemental Figure 4.
- **Data for Supplemental Figure 5 – HNH Inteins Aligned.fna** Nucleotide sequence alignment of identified HNH-containing inteins, used to generate phylogeny in Supplemental Figure 5.
- **Data for Supplemental Figure 6 – Anti-Sense LAGLIDADG Inteins Aligned.fna** Nucleotide sequence alignment of identified anti-sense LAGLIDADG-containing inteins, used to generate phylogeny in Supplemental Figure 6.
- **Supplemental Invader Sim Results Graphs.zip** Summary plots of results from searching the Invader-Sim simulated mixed intein and non-intein containing sequence databases to test ICE-BLAST.

### Supplemental Methods Information

#### ICE-BLAST: Detailed Methodology

ICE-BLAST loops through the following steps (described visually in Figure 1 - A). First, the input sequence is used as the query in a PSI-BLAST (v 2.12.0+) (Altschul et al. 1997) versus a training database known to contain sequences related to the query (but it is not the database of interest, e.g. Uniref50 (Suzek et al. 2015)). The resulting position specific scoring matrix (PSSM) is then used to run a second PSI-BLAST search against the protein database of interest (out-database in Figure 1 - A). The resulting matches from this second search are filtered based on e-value, percent identity, and query coverage, after which the filtered matches are saved into memory. These filtered matches are clustered using USEARCH (v11.0.667) (Edgar 2010) at the percent sequence identity specified by the user (default 70%) using cluster_fast mode. Finally, the centroids of these sequence clusters are used as the query sequences for the next iteration of the algorithm. This loop continues until (a) no new matches are found in the second PSI-BLAST search, (b) the clustering step produces no new centroid sequences, or (c) the maximum number of iterations (specified by the user) is reached.

ICE-BLAST also has a domain specific mode which can be enabled by the user; this mode treats the query sequence as a sub-sequence of any subject proteins. The algorithm proceeds through the same loop as before, but instead of using the entire protein match from the second PSI-BLAST search, only the portion matching the query is used for clustering. Importantly, this match is confined to an upper and lower bound set by the user when invoking domain specific mode. They are equivalent to the average length of the query sequence(s) plus or minus the user specified percentage of the average length (i.e. an average sequence length of 100aa and a domain specific mode cutoff of 20% would lead to an upper bound of 120aa and a lower bound of 80aa).

**Invader-Sim: Detailed Methodology**

Invader-Sim simulates phylogenies of inteins and exteins using PAML-EVOLVER (Yang 2007). Invader Sim then pairs these phylogenies 1-1 at random such that each intein tree is paired with one extein tree. Intein invasion is simulated by randomly selecting one tip from the intein phylogeny to "invade" a randomly selected tip from the extein phylogeny. The intein tip with the closest patristic distance in the intein phylogeny to the previous intein tip is chosen as the next invading sequence. The target of this invasion is chosen by a Markov chain using an exponential distribution and two uniform random variables for selection and rejection. This process continues until all inteins have “invaded” an extein sequence and have been paired. Note that this need not mean all extein sequences have been invaded. For testing purposes some extein sequences were always left unpaired (see “Parameters Used In Testing” section below for more detail).

Once the simulated invasion process is complete, amino acid sequences are simulated for the intein and extein phylogenies using PAML-EVOLVER. Intein sequences are inserted into their paired extein counterpart at an insertion site randomly selected within the middle two quartiles of the extein's amino acid sequence (this position is used for all invading inteins within the current extein-intein phylogeny pair). This process is repeated for every sample the user requests.

A modification was made for a second test such that only a subsample of the intein tips simulated are used to invade the extein sequences (to provide a more realistic database with only partial sampling of the true intein population). This subsample of intein sequences (specified by the user) is chosen at random from the entire intein phylogeny, and only those randomly selected sequences will be used in the subsequent steps of the algorithm. All other tips in the intein phylogeny are ignored in regards to selecting the closest tip during the invasion process, and are not used to simulate sequences.

**Invader-Sim: Study Parameters**

Invader-Sim simulations in this manuscript used the following parameters. The extein phylogenies were simulated with a birth rate of 1, a death rate of 1, 1000 tips, a sampling fraction of 0.1, and a mutation rate of 0.1. The intein phylogenies were simulated with a birth rate of 2, a death rate of 1, a sample fraction of 0.5 and a mutation rate of 1. The number of tips for the intein phylogenies differed between the sub-sample and original methodology for dataset construction. In the latter we simulated a phylogeny with 100 intein tips, whereas the former simulated a phylogeny with 1000 intein tips but randomly selected 100 of said tips for invasion.

Intein invasion into exteins was simulated with a window size of 100, and an exponential probability distribution with a lambda of 0.5.

Extein amino acid sequences were simulated to be 500 amino acids in length, using an alpha parameter of 0.5, 4 rate categories, and a proportional model. Intein sequences were simulated with a length of 300 amino acids, an alpha parameter of 1, 4 rate categories, and a proportional model.

### Supplemental Results and Discussion

#### ICE-BLAST Testing using Simulated Data

For each simulated dataset, we simulated the evolutionary histories of 1000 homologous extein sequences, and 100 homologous intein sequences. These inteins were invaded at random into a random subset of the extein sequences, and amino acid sequences were then derived from the simulated phylogenies. For each simulation, we took 1 random simulated intein to act as the query, and tested true positive and false positive recovery rates. We conducted tests not only using ICE-BLAST, but also with standalone PSI-BLAST, and standalone BLASTP, in order to provide a relative measure of the program's false positive and true positive rates.

We ran 100 simulated datasets as described above, and then searched this dataset using each of the programs listed above, while varying the e-value cutoff of an acceptable match (for all 3), and the percent identity used for clustering between searches (ICE-BLAST only). The total number of matches, true positive matches, false positive matches, and the average length of the match was recorded in Supplemental Table 4 (data within this table is broken down by experiment).

We found that none of the runs recovered false positive matches, and that both PSI-BLAST and ICE-BLAST almost always recovered 100 out of 100 simulated intein sequences. BLASTP, as expected, did not perform well at higher e-value cutoff. This is especially true of e-values below E-15, at which point BLASTP sometimes recovered less than 90% of the simulated inteins.

We also ran simulations as described prior, but with a modification to the intein simulation process. In an effort to simulate our incomplete sampling of inteins in normal databases, we conducted 100 simulations where we simulated 1000 intein sequences, but only used 100 for the invasion process. This “sub-sample” dataset was then used in the same manner as the experiment described prior, and with similar results (see Supplemental Table 4). There were no significant differences between the results of these two simulations.

Finally, we used the 100 samples from the original simulations, and the 100 samples from the sub-sample simulations to create two unified datasets consisting of all intein and extein sequences from their class of simulation. These two datasets (the “unified sub-sample” and “unified original”) were similarly tested on using the 3 programs previously mentioned. Again, there was no significant difference between the results of these experiments and the two described prior; with one exception. BLASTP performed better in the unified dataset experiment, with most of the poor performance outliers disappearing.

Overall the results of our simulation experiments can be distilled down into 2 key findings. First, the ICE-BLAST performs comparably to PSI-BLAST in regards to it false positive rate; a positive result, considering that the primary concern of this iterative clustering approach was a high false positive rate. Second, we found that, on average, PSI-BLAST and ICE-BLAST recovered 4 to 5 more residues of the simulated intein than BLASTP. However, the primary driver of recovery length was ultimately the particularities of each simulated samples. Outliers for intein length recovery for all three methods came from the same simulated samples.

Summary statistics for each of the different types of experiment (the baseline, the 100 tip subsample, all baseline datasets unified, and all 100 tip subsample datasets unified) can be found in Supplemental_Material_Invader_Sim_Results_Graphs.zip. These figures compare true positive rates and the average length of the retrieved simulated intein sequence against varying e-value and percent identity cutoff values (for BLAST and uclust respectively).

**Intein Insertion Sites Analysis**

The numeric position for intein insertion within its host extein within our dataset varied significantly even in a single related cluster of inteins; however, this variation was largely the result of differences in host extein architecture, and not a difference in the specific insertion sequence being targeted. As a result, we chose to include the target DNA sequence (+/-30nt) from the target site instead of a numeric position within the amino acid sequence. Given the composition of our dataset, we found consensus sequences constructed from all insertion sequences heavily biased the consensus to the inteins within terminase genes. To combat this bias we created consensus sequences (at 60% identity) of the insertion sequences for each of our intein clusters. We then grouped these together based on the type of homing endonuclease contained within the intein, as described above (excluding the mini-inteins). A consensus sequence was constructed from said consensus sequences (Supplemental Table 5, Supplemental Figure 12).

The consensus insertion site for in-frame LAGLIDADG HEN containing inteins was dominated by the much more prevalent sequences from class 1 inteins. However, if class 3 inteins were separated from those class 1 inteins, we found a significantly more resolved insertion site consensus. Almost all of the class 3 inteins found in our dataset seem to require an adenine at the -1 position, and have more stringent +1 requirements when compared to the class 1 inteins whose consensus sequence has higher conservation on the +2 and +3 residues.

The VSR-like HEN containing intein insertion site consensus sequence does not quite match the expected target site for VSR proteins ( CT(AT)GN or NT(AT)GG ); however, since this VSR-like HEN is divergent compared to known VSR HENs, it is not unlikely that it would target a slightly different nucleotide motif.

We could not draw significant observations from the consensus sequences of the anti-sense LAGLIDADG HEN containing inteins, the HNH HEN containing inteins, nor the putative endonuclease VII-like HEN containing inteins. The anti-sense LAGLIDADG HEN containing inteins, and endonuclease VII-like HEN containing inteins, invaded closely related extein sequences, and thus provided a very uniform and uninformative consensus sequence for their insertion site. The HNH HEN containing inteins proved to be a diametrically opposed case, where the inteins within our dataset had invaded sufficiently divergent extein sequences such that a consensus sequence with any degree of meaningful information quality could not be constructed.

**Sequence Conservation in Inteins and Exteins**

The conflicts between intein and extein phylogenies reveal that inteins are frequently transferred between genes. This and the often-sporadic distribution of inteins on the extein phylogeny also reveals that in most instances inteins are frequently lost and gained. Here we show that for divergent extein/intein pairs inteins are less conserved than the exteins, and that the one intein/extein family in actinobacteriophages for which reliable codon-based sequence alignments could be calculated the intein sequence contained fewer sites under strong purifying selection than the extein sequences.

Given the frequent transfer of inteins between extein sequences, it is difficult to assess the relative sequence conservation of inteins and exteins. Pairwise sequence comparisons of homologous genes with divergent inteins inserted in the same location in the extein illustrate that inteins can diverge much faster than the extein sequences (Supplementary Fig.13). However, the selection of intein containing genes obviously biases the result. If homologous genes with nearly identical inteins are selected, the inteins appear more conserved than the exteins.

We argue that the finding of identical intein sequences reflects their recent transfer between exteins.  An alternative explanation for finding identical inteins in divergent extein sequences could be that the inteins are under stronger selection for sequence conservation than the extein sequences, although the absence of even synonymous substitutions in the intein sequences, argues that these identical intein sequences diverged recently and did not coevolve with the extein sequences. Phylogenetic reconstruction and estimation of relative divergence rates and selection pressures is difficult because sequences within intein clusters (as defined here, see Supplemental Table 1) are often nearly identical, and between intein clusters so divergent that the sequences cannot be aligned with confidence.  For example, inteins inserted into the same site of the minor capsid proteins have even acquired different homing endonucleases located in different reading frames.

An exception are the nucleotidyltransferases (intein clusters #1 and #69).  The amino acid sequence alignment via muscle (Edgar 2004) as implemented in seaview (Gouy et al. 2021) introduces only two single amino acid gaps in the intein from cluster#1 . Nucleotide sequences for the inteins of clusters #1 and #69 can be found in Supplemental Data in Identified Inteins – Intein Nucleotide Sequences.fna.

Analyzing the nucleotide sequence alignments based on this amino acid sequence alignment in MrBayes (lset nst=6 rates=gamma; mcmc ngen=10000000;)

(Ronquist et al. 2012) in separate analyses for the intein and extein sequences estimated the alpha shape parameter for the gamma distribution for the exteins to 0.0654 with a 95% High Probability Density (HPD) interval of [0.0126, 0.0946], reflecting very strong selection acting on most sites in the extein. In contrast, the shape parameter for the intein sequences was 4.7 (HDP interval [1.9202, 8.0112]) suggesting a much less extreme among site rate variation whose distribution is similar to a normal distribution, with only few sites within the intein sequence under strong purifying selection.

Similarly, an analysis of the nucleotidyltransferase datasets using a codon model (Nielsen and Yang 1998) that allows for different classes for the ratios for the rates of non-synonymous (dN) to synonymous (dS) substations (MrBayes settings: *lset nst=2 rates=gamma nucmodel=codon omegavar=Ny98;* mcmc ngen=200;000; burnin=50,000) reveals fewer sites under purifying selection in case of intein, and the dN/dS rate estimated for the intein is higher than for the extein, i.e., the intein has fewer sites under purifying selection and the average strength of selection acting on these sites lower in case of the intein. A summary of the estimated parameters describing the strength of selection acting in the nucleotidyltransferase inteins and exteins is given in Supplemental Table 6.

These analyses may have been impacted by alignment ambiguities; however, the findings that the intein has fewer sites under purifying selection and that the selection pressure on these sites is weaker than for the extein agree with unpublished results (Gogarten) for the intein in the vacuolar ATPase catalytic subunit in yeast, for which many more sequences that can be aligned with confidence are available.

### Supplemental Material References


Altschul SF, Madden TL, Schäffer AA, Zhang J, Zhang Z, Miller W, Lipman DJ. 1997. Gapped BLAST and PSI-BLAST: a new generation of protein database search programs. *Nucleic Acids Research* 25:3389–3402.

Edgar RC. 2004. MUSCLE: multiple sequence alignment with high accuracy and high throughput. *Nucleic Acids Res* 32:1792–1797.

Edgar RC. 2010. Search and clustering orders of magnitude faster than BLAST. *Bioinformatics* 26:2460–2461.

Gouy M, Tannier E, Comte N, Parsons DP. 2021. Seaview Version 5: A Multiplatform Software for Multiple Sequence Alignment, Molecular Phylogenetic Analyses, and Tree Reconciliation. *Methods Mol Biol* 2231:241–260.

Madeira F, Madhusoodanan N, Lee J, Eusebi A, Niewielska A, Tivey ARN, Lopez R, Butcher S. 2024. The EMBL-EBI Job Dispatcher sequence analysis tools framework in 2024. *Nucleic Acids Res* 52:W521–W525.

Nielsen R, Yang Z. 1998. Likelihood models for detecting positively selected amino acid sites and applications to the HIV-1 envelope gene. *Genetics* [Internet] 148:929–936. Available from: http://www.ncbi.nlm.nih.gov/entrez/query.fcgi?cmd=Retrieve&db=PubMed&dopt=Citation&list_uids=9539414

Ronquist F, Teslenko M, van der Mark P, Ayres DL, Darling A, Höhna S, Larget B, Liu L, Suchard MA, Huelsenbeck JP. 2012. MrBayes 3.2: Efficient Bayesian Phylogenetic Inference and Model Choice Across a Large Model Space. *Syst Biol* [Internet] 61:539–542. Available from: https://www.ncbi.nlm.nih.gov/pmc/articles/PMC3329765/

Suzek BE, Wang Y, Huang H, McGarvey PB, Wu CH, the UniProt Consortium. 2015. UniRef clusters: a comprehensive and scalable alternative for improving sequence similarity searches. *Bioinformatics* 31:926–932.

Yang Z. 2007. PAML 4: Phylogenetic Analysis by Maximum Likelihood. *Molecular Biology and Evolution* 24:1586–1591.
